# Conformational rearrangements upon start codon recognition in human 48S translation initiation complex

**DOI:** 10.1101/2021.12.17.473132

**Authors:** Sung-Hui Yi, Valentyn Petrychenko, Jan Erik Schliep, Akanksha Goyal, Andreas Linden, Ashwin Chari, Henning Urlaub, Holger Stark, Marina Rodnina, Sarah Adio, Niels Fischer

## Abstract

Selection of the translation start codon is a key step during protein synthesis in human cells. We obtained cryo-EM structures of human 48S initiation complexes and characterized the intermediates of codon recognition by kinetic methods using eIF1A as a reporter. Both approaches capture two distinct ribosome populations formed on an mRNA with a cognate AUG codon in the presence of eIF1, eIF1A, eIF2–GTP–Met-tRNA_i_^Met^, and eIF3. The ‘open’ 40S subunit conformation differs from the human 48S scanning complex and represents an intermediate preceding the codon recognition step. The ‘closed’ form is similar to reported structures of complexes from yeast and mammals formed upon codon recognition, except for the orientation of eIF1A, which is unique in our structure. Kinetic experiments show how various initiation factors mediate the population distribution of open and closed conformations until 60S subunit docking. Our results provide insights into the timing and structure of human translation initiation intermediates and suggest the differences in the mechanisms of start codon selection between mammals and yeast.

## Introduction

In eukaryotes, mRNA is recruited to the 43S pre-initiation complex, 43S PIC, which consists of the 40S ribosomal subunit, translation initiation factors eIF1, eIF1A, eIF3, eIF5, and a ternary complex (TC) composed of eIF2, GTP, and Met-tRNA_i_^Met^. 43S PIC binds to the 5’ end of the mRNA and scans along the 5’ untranslated region (5’UTR) in the 3’ direction to find the start codon (AUG) within the context of an optimal Kozak sequence. Start codon recognition stabilizes the 48S initiation complex (48S IC), initiates dissociation of eIF1, eIF1A, eIF2, and eIF5, and promotes recruitment of the 60S ribosomal subunit to form 80S IC ready to enter the elongation cycle of protein synthesis. Start codon selection establishes the open reading frame and determines the amino acid sequence of the synthesized protein. The frequency of translation initiation at a given AUG start codon defines translational efficiency of the mRNA, which shapes the composition of the cellular proteome. Compromised fidelity of AUG selection is a common feature of human diseases such as neurodegeneration or cancer (Costa-Mattioli and Walter, 2020; Moon and Parker, 2018; Robichaud et al., 2019).

Translation initiation factors play important roles in mRNA recruitment and start codon selection. eIF1A binds to the A site of the 40S subunit. In yeast, mutations of eIF1A and eIF1 influence the fidelity of the start codon selection *in vivo* and *in vitro* (Fekete et al., 2007; Maag et al., 2006; Nanda et al., 2009; Nanda et al., 2013). Mutations of eIF1A N-terminal tail (NTT) enhance leaky scanning, whereby the scanning ribosomes bypass the first AUG start codon and initiate translation at downstream AUG start codons (Fekete et al., 2007). Mutations of eIF1A C-terminal tail (CTT) have an opposite (inhibitory) effect on the leaky scanning and enhance initiation at a near-cognate UUG codon within a 5’-proximal Kozak sequence. Mutations of eIF1 that change its dissociation rate from the 48S PIC affect the selection of cognate- and near-cognate codons *in vitro* (Nanda et al., 2013). Premature release of eIF1 promotes translation initiation at sub-optimal translation initiation sites (Cheung et al., 2007; Nanda et al., 2009).

In both east and mammals, eIF1, eIF1A, and (in mammals) eIF3 enhance mRNA scanning by inducing an open conformation of the mRNA channel of the 40S subunit and by coordinating the TC binding (Brito Querido et al., 2020; des Georges et al., 2015; Hashem et al., 2013a; Hashem et al., 2013b; Kumar et al., 2016; Lomakin and Steitz, 2013; Obayashi et al., 2017; Passmore et al., 2007; Weisser et al., 2013). eIF1, which binds to the P site of 40S subunit, interferes with the accommodation of Met-tRNA_i_^Met^ in the P site during scanning. Upon codon recognition, eIF1 is released from the ribosome. In mammals, eIF1 and eIF1A act synergistically to mediate the assembly of h48S IC on the initiation codon and enhance binding affinities of each other to the 40S subunit (Pestova et al., 1998; Sokabe and Fraser, 2014). eIF1 inhibits premature GTP hydrolysis by eIF2 until an AUG codon is recognized (Brito Querido et al., 2020; Lomakin and Steitz, 2013; Unbehaun et al., 2004). In the absence of eIF1 or eIF3, h48S IC does not scan mRNA and remains in the proximity of the mRNA 5’-end (Kumar et al., 2016). In the late stage of h48S IC formation, eIF1A competes with eIF5 for binding to eIF5B, allowing dissociation of eIF5-eIF2 complex after GTP hydrolysis by eIF2 (Lin et al., 2018).

eIF5 is a GTPase-activating protein (GAP) of eIF2. In yeast, eIF5 binds to eIF2 during scanning and induces GTP hydrolysis by eIF2, but the reaction product, inorganic phosphate (Pi) remains bound until the ribosome recognizes the start codon, which triggers Pi release and the subsequent eIF2 dissociation (Algire et al., 2005; Algire et al., 2002; Llacer et al., 2018). By displacing eIF1, eIF5 facilitates the accommodation of Met-tRNA_i_^Met^ and enables gated release of Pi from eIF2, which effectively ends the scanning process (Algire et al., 2005; Llacer et al., 2018; Luna et al., 2013; Luna et al., 2012; Saini et al., 2014). This is different in mammals, where GTP hydrolysis by eIF2 is induced upon codon recognition by relieving inhibition by eIF1 (Pestova and Kolupaeva, 2002; Pisareva and Pisarev, 2014; Unbehaun et al., 2004). Finally, eIF5B is a translational GTPase that facilitates 60S subunit docking (Acker et al., 2009; Pestova et al., 2000; Pisareva et al., 2007; Wang et al., 2019). Recruitment of 60S subunit induces dissociation of the remaining initiation factors and marks the onset of translation elongation.

Structural and biochemical work suggests that the 48S complexes adopt different conformations during scanning and upon start codon recognition. The structure of yeast 48S IC trapped on a near-cognate AUC instead of the AUG codon in the mRNA (y48S AUC) shows an ‘open’ 40S conformation with Met-tRNA_i_^Met^ in the P_out_ state (Hussain et al., 2014; Llacer et al., 2015). In this intermediate, scanning is stopped by partial base pairing of Met-tRNA_i_^Met^ with the AUC codon. AUG recognition induces the accommodation of Met-tRNA_i_^Met^ into the P site (P_in_ state) and tightens the mRNA binding channel around the anticodon stem, forming a ‘closed’ state of the y48S AUG (Hussain et al., 2014; Llacer et al., 2015; Llacer et al., 2018). This conformation is stabilized by contacts between the NTT of eIF1A with the tRNA-mRNA duplex (Hussain et al., 2014; Llacer et al., 2015; Llacer et al., 2018). AUG recognition induces the release of eIF1 and eIF2β from the P site, Pi release from eIF2, and reorganization of eIF3 (Algire et al., 2005; Kapp and Lorsch, 2004; Llacer et al., 2015; Llacer et al., 2018; Maag et al., 2005). The existing structures of mammalian 48S IC assembled on AUG are similar to the closed y48S AUG (Eliseev et al., 2018; Simonetti et al., 2020). Recent structural work has captured human 48S complex in the course of mRNA scanning (Brito Querido et al., 2020), which shows a distinct 40S subunit conformation and no codon-anticodon recognition in the P site. One important difference between yeast and mammalian 48S IC assembly concerns the role of eIF3 in AUG recognition. Yeast eIF3 appears to be not essential (Maag et al., 2006), whereas mammalian eIF3 is indispensable for h43S PIC and h48S IC formation (Kumar et al., 2016; Pestova and Kolupaeva, 2002; Sokabe and Fraser, 2014; Sokabe et al., 2012). While high-resolution structures and detailed biochemical studies of y48S IC provide important insights into the mechanism of start codon selection in yeast, structural view on the process in higher eukaryotes is just starting to emerge. The kinetic data on translation initiation in mammals are scarce compared to the detailed analysis available for the yeast system (Algire et al., 2005; Kapp and Lorsch, 2004; Llacer et al., 2015; Llacer et al., 2018; Maag et al., 2005), which calls for time-resolved studies in the mammalian system.

Here we use a fully reconstituted *in vitro* translation system from human cells to study the assembly of human 48S IC by single-particle cryo-EM and rapid kinetic approaches. Our cryo-EM data show two distinct conformations of the human 48S complex assembled on an mRNA with a cognate AUG start codon, prior to and after codon-anticodon recognition. Although both complexes contain the same set of initiation factors, e.g. eIF1A, eIF1, TC, and eIF3, their arrangement at the decoding center and the conformations of the 40S subunit are different. Comparison to previously reported structures, in particular the open and closed y48S, as well as h48S scanning complexes, suggests that the open h48S AUG may represent an early intermediate of the codon recognition process. Kinetic analysis using eIF1A dissociation as a diagnostic assay for changes in ribosome conformations suggests how eIFs and start codon recognition remodel the ribosome from 43S PIC and 48S PIC to 48S IC and finally, how eIF5B remodels 48S IC conformation for the 60S subunit joining. This work provides further evidence for the conserved and distinct features of start codon-recognition mechanism in mammals and yeast.

## Results

### Structures of h48S AUG in the open and closed state

To analyze 48S IC assembled on the AUG start codon by cryo-EM, we reconstituted complexes using purified human 40S subunits, eIF1, eIF1A, eIF2–GTP–Met-tRNA_i_^Met^ (TC), eIF3, in the presence of eIF4A and 4B, and *in-vitro* transcribed mRNA (Figure 1 – figure supplement 1 and Methods) (Pestova et al., 1998; Pestova and Kolupaeva, 2002; Pisarev et al., 2007). The model mRNA (Figure 1 – figure supplement 2A) has an unstructured 5’UTR, which overcomes the requirement for 5’ capping and eIF4F recognition, followed by an optimal Kozak sequence, a short open reading frame coding for a peptide MVRFKA, and a 3’UTR sequence that serves as a primer binding site for the primer-extension inhibition (toe-printing) assay (Pestova and Kolupaeva, 2002). Functional activity of 48S and 80S complexes was verified by toe-printing (Figure 1 – figure supplement 2A). In mammals, eIF1, eIF1A, TC and eIF3 are the minimal set of essential factors to reconstitute the 48S IC on an unstructured and uncapped mRNA (Pestova et al., 1998; Pestova and Kolupaeva, 2002) (Figure 1 – figure supplement 2B). 48S IC causes a strong toe-printing stop on the cognate start codon (AUG), which is not observed with the non-cognate codon (CUC), while on the near-cognate codon (AUC) 48S IC forms a specific, but labile complex (Figure 1 – figure supplement 2C). These human 48S initiation complexes formed in the presence of eIF1A, eIF1, TC, eIF3 but in the absence of eIF5 or eIF5B are denoted in the following as h48S AUG, h48S CUC and h48S AUC.

For the cryo-EM work on h48S AUG complexes, we used the GraFix procedure (Kastner et al., 2008) to stabilize the complexes by crosslinking with bis(sulfosuccinimidyl)suberate (BS3), p-maleimidophenyl isocyanate and glutaraldehyde. After sorting cryo-EM images according to the 40S subunit conformation and the presence of eIFs (Figure 1 – figure supplement 3 and Methods), we obtained two final cryo-EM reconstructions depicting h48S AUG in an open and closed state, respectively (Figure 1 and Supplementary table 1). In both states, we were able to model almost all components of the complex, except for eIF1 and the C-terminal domain of eIF2β in the closed state (see below). In the final structures, the occupancy of 40S subunits with eIF3 was high ≥80% (estimated by *in silico* sorting of cryo-EM data, see Methods for details) and almost all 13 subunits of eIF3 could be placed, except for the flexible peripheral subunit eIF3i (Brito Querido et al., 2020) and the loosely associated subunit eIF3j (Fraser et al., 2007; Sokabe and Fraser, 2017). Analysis of the h48S AUG complexes using crosslinking mass spectrometry (XLMS) supported our structural models (Figure 1 - figure supplement 4, Supplementary tables 2-5 and Methods). 91% of the crosslinks fall into the 30 Å range for both the crosslinkers used (BS3 and LC-SDA). Crosslinks that exceed the distance restrictions (35-45 Å) can be explained by local flexibility and dynamic regions of the structures. We note that more crosslinks could be mapped onto the open state, especially because eIF1 and some parts of eIF3c could only be modeled in the open state (see below).

**Figure 1.**
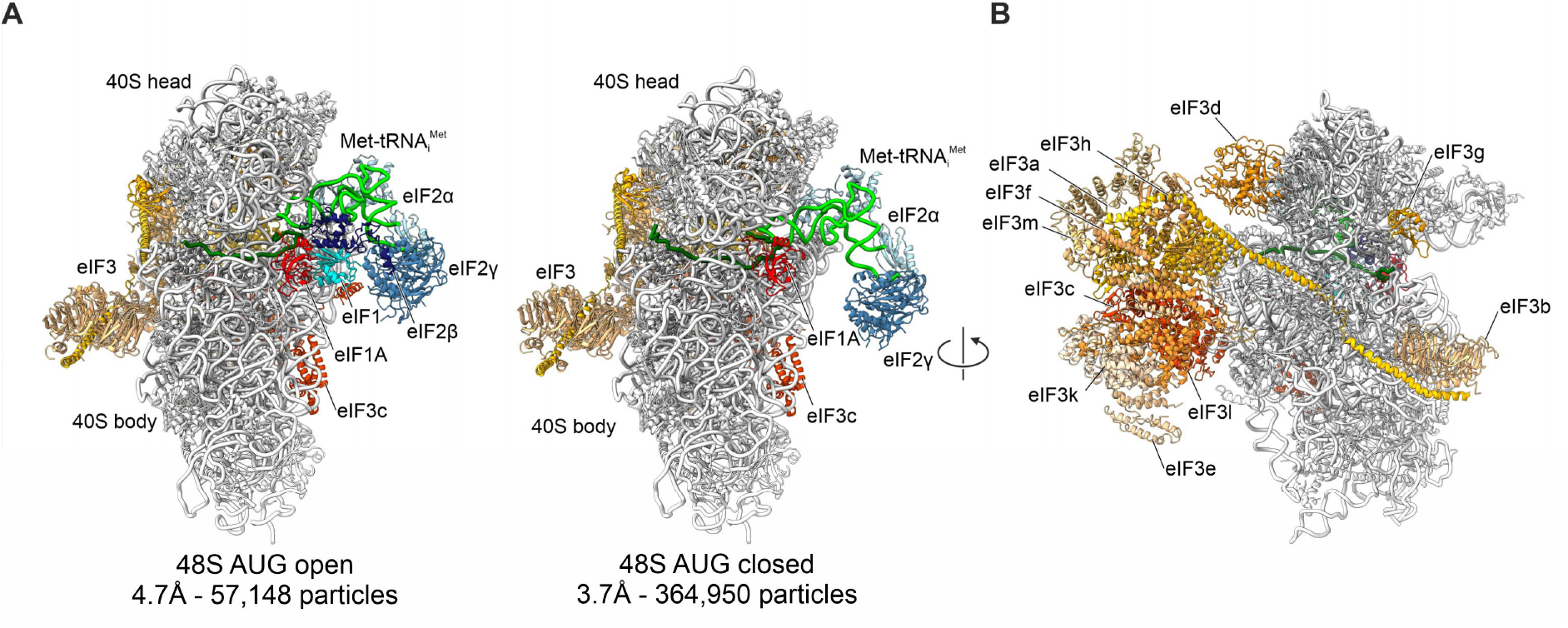
Structures of the h48S AUG complex in open and closed states. **(A)** Overall structures of the open (left) and closed (right) states assembled on mRNA with cognate AUG start codon. Structures are shown as ribbon models with the respective resolution and particle numbers used for the final cryo-EM maps. In the closed state, the density for eIF1 and eIF2β is missing. **(B)** Structural assignment of eIF3 subunits in h48S AUG closed. All subunits of eIF3 could be placed except for eIF3i and eIF3j. Individual subunits of eIF3 are color-coded as indicated.

The overall conformation of the 40S subunit is clearly different in the open and closed states (Figure 2A and Figure 2 – figure supplement 1). The 4.7 Å cryo-EM structure of the h48S AUG open represents a minor population of the complexes (~16%, Figure 1) that do not appear to form the codon-anticodon interactions Figure 2B), although the mRNA contains the AUG codon and all the initiation components are present (Figure 1). In our open state, the 40S head domain is tilted towards the solvent site, which opens the decoding center (Figure 2A and Figure 2 – figure supplement 1A) and the mRNA entry latch (Figure 2C). Probably due to the absence of codon-anticodon interaction, only the mRNA backbone is resolved in h48S AUG open; the lack of information for the bases prevents identification of the codon in the P site. Met-tRNA_i_^Met^ is in the P_out_ conformation and contacts the 40S head domain. Specifically, two conserved GC base pairs in the tRNA anticodon stem interact with G1639 to A1641 in helix 29 (h29) of 18S rRNA mostly via unspecific RNA-RNA backbone interactions (Figure 2C,D), similarly to the interactions observed in yeast (Llacer et al., 2015). eIF2α and eIF2β reach towards the decoding center and interact with the tRNA, whereas eIF2γ points away from the 40S subunit body (Figure 2C). eIF1 binds close to the P-site codon, while the C-terminal domain (CTD) of eIF2β interacts with eIF1 and contacts the anticodon-stem loop of the tRNA, thereby stabilizing the tRNA P_out_ conformation (left panels in Figure 2B,D). eIF1A binds to the A site of the decoding center between h18 and h44 of 18S rRNA and proteins uS12 and eS30 of the 40S body domain (Figure 2E,F and Figure 2 – figure supplement 1B), as in all reported 48S IC structures (Brito Querido et al., 2020; Llacer et al., 2015; Llacer et al., 2018; Simonetti et al., 2020). An α-helical element (residues 265-278) in the N-terminal domain (NTD) of eIF3c, which is specific for human eIF3, interacts with eIF1 in a similar way as in the h48S·scan complex (Figure 2C and Figure 2 – figure supplement 1C; (Brito Querido et al., 2020)).

**Figure 2.**
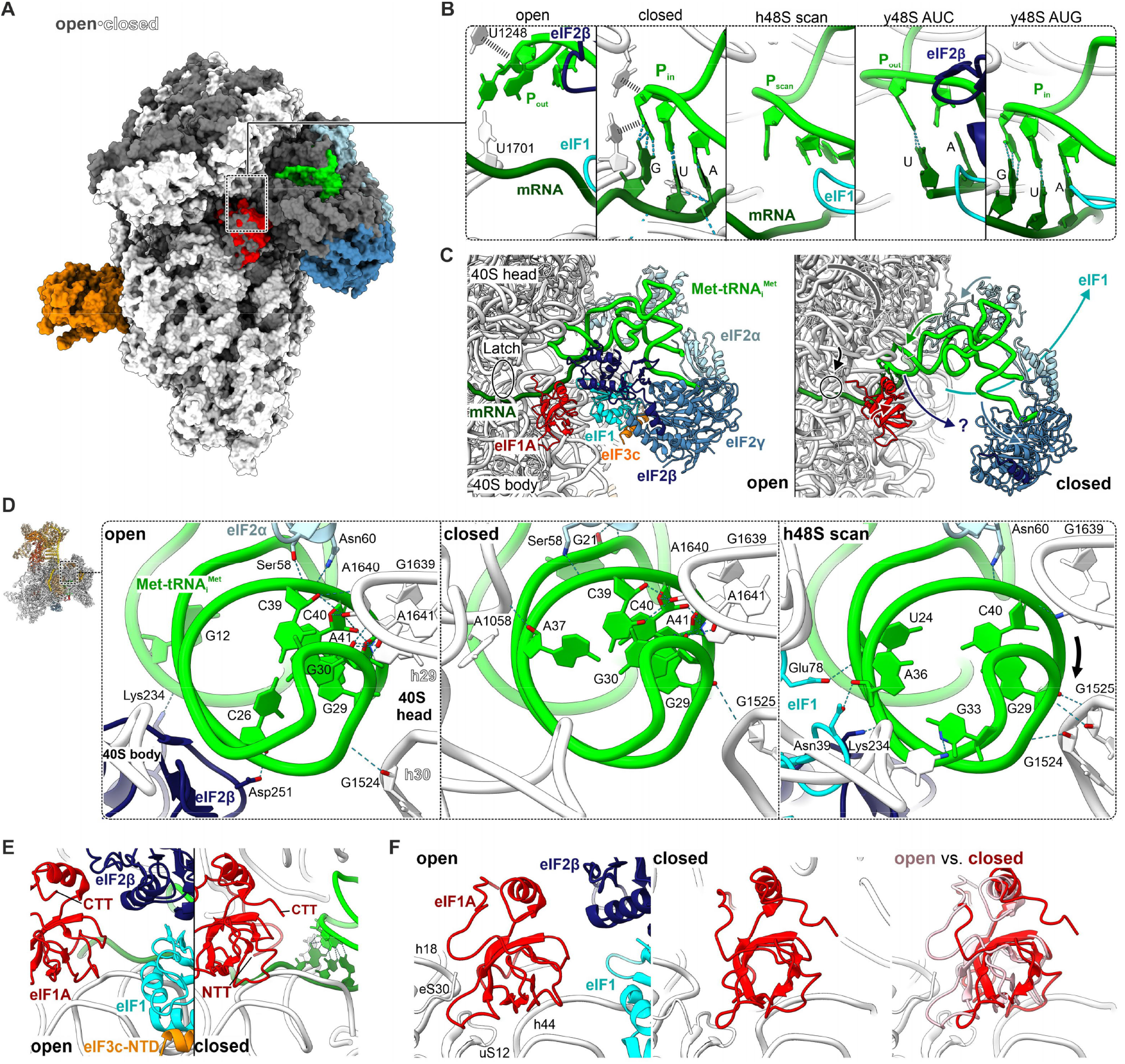
Two distinct conformations of the h48S AUG complex. **(A)** Superposition of the h48AUG in the open state (all components of the complex are shown in grey) and the closed state (40S subunit, white; eIFs colored as in Fig. 1: eIF1A, red; Met-tRNA_i_^Met^, green; eIF2α, light blue; eIF2γ, blue; eIF3, orange); atomic models for both complexes are shown in space fill representation. Here and in other Figures, structures were aligned by the 40S body domain, if not mentioned otherwise. **(B)** Close-ups of the P site in 48S IC complexes from human and yeast. Open, closed: h48S AUG states described here. h48S scan: human 48S IC with non-cognate codon CUC replacing the AUG codon (PDB 6ZMW, (Brito Querido et al., 2020)). y48S·AUC: yeast 48S IC open state with a near-cognate AUC codon (PDB 3JAP, (Llacer et al., 2015)). y48S·AUG: yeast 48S IC with cognate AUG and eIF1 bound in closed state (PDB 3JAQ, (Llacer et al., 2015)). Note the repositioning of both Met-tRNA_i_^Met^ from P_out_/P_scan_ to P_in_ and mRNA upon 40S head domain closure resulting in codon-anticodon interaction. **(C)** Rearrangements of the TC in the open vs. closed h48S AUG. Arrows mark major conformational changes upon 40S head (grey) movement from open to closed state: closure of the mRNA entry latch (black circle); movement of Met-tRNA_i_^Met^ (green) and eIF2α (light blue) towards and of eIF2γ away from the 40S subunit; dissociation of eIF1 (cyan) and movement of eIF2β (purple) away from the P site; and repositioning of eIF1A (red). **(D)** Interactions between the ASL of Met-tRNA^Met^ (green) with the 40S head domain (18S rRNA, white). Note that tRNA contacts in the h48S AUG open and closed states are similar, but different in h48S scan state (PDB 6ZMW (Brito Querido et al., 2020)). Dashes indicate residues that are within H-bond distance. Arrow marks the shift in interactions from h29 to h30. Structures were aligned by the ASL of initiator tRNA. **(E)** Close-ups of the decoding center. Upon 40S head closure, eIF1 and eIF2 move out of the decoding center, while eIF1A and its N-terminal tail (NTT) move towards the P site. Note the α-helix in the NTD of eIF3c (eIF3c-NTD), which interacts with eIF1 in the open state and is not resolved in the closed state. **(F)** Repositioning of eIF1A. Position and interactions of eIF1A in the open (left) and closed states (center), and superposition (right) of eIF1A in open (light red) and closed state (red). uS12 and eS30, proteins of the 40S subunit; h44, helix 44 of 18S rRNA.

The major population of h48S AUG (~84%) adopts a closed state, denoted as h48S AUG closed. The structure, obtained at 3.7 Å resolution, shows the closure of the 40S head domain and of the mRNA entry latch upon start codon recognition (Figure 2A-D and Figure 2 – figure supplement 1A. Met-tRNA_i_^Met^ moves into the P site (P_in_) and the mRNA changes its path to base pair with the tRNA (Figure 2B). The TC, and in particular tRNA_i_^Met^, does not change its interactions with the 40S head domain, but moves together with it upon 40S subunit closure (Figure 2D). eIF1 and the CTD of eIF2β appear to move out of the decoding center, as the respective density is not traceable in the closed state (Figure 2C). Upon release of eIF2β from the decoding center, the peripheral part of the TC, in particular eIF2α and the acceptor stem of Met-tRNA_i_^Met^, become more flexible and are not as well defined as in the open state. In the closed state there is also no density for the α-helical element of eIF3c, which appears to dissociate together with eIF1 from the decoding center upon start codon recognition. Moreover, formation of the closed state induces a rearrangement of eIF1A, which now lacks the contacts with h18 and eS30 and moves closer to the P site where its N-terminal tail reaches towards the tRNA (Figure 2E,F and Figure 2 – figure supplement 1B).

### Comparison to reported 48S IC structures

The h48S AUG open state, which was not captured in other 48S IC structures with AUG, is similar to the near-cognate y48S AUC complex (Llacer et al., 2015)(Figure 3A). The main difference is in mRNA-tRNA interactions, as in the y48S AUC complex the anticodon of tRNA_i_^Met^ interacts with the U of the near-cognate AUC codon (Figure 2B), whereas in h48S AUG open complex the mRNA takes a different path and appears too far away from tRNA_i_^Met^ for base pairing. The present h48S AUG closed state resembles previous high-resolution structures of 48S IC obtained with a cognate AUG codon, both from yeast (y48S AUG, (Llacer et al., 2015; Llacer et al., 2018)) and rabbit (r48S AUG, assembled on β-globin mRNA, (Simonetti et al., 2020)), except for the orientation of eIF1A (see below) (Figure 3A and Figure 3 – figure supplement 1A). Notably, closed y48S AUG contains eIF5, which replaces eIF1 and appears to stabilize eIF2β and eIF2α (Llacer et al., 2018) similarly to eIF1 in the h48S AUG open state. In y48S AUG, two slightly distinct positions of eIF2γ were identified that differ from the present h48S AUG closed state by reaching further towards the decoding center, most likely due to the interactions of eIF5 with the decoding center in the yeast complexes (Llacer et al., 2015; Llacer et al., 2018). Comparing r48S AUG with the h48S AUG closed complex, the similarities are in the overall 40S subunit conformation and the positions of Met-tRNA_i_^Met^. In both structures the density for eIF1, eIF2β, eIF3i and eIF3j are not resolved. However, there are also differences, for example, a somewhat different orientation of eIF2y, eIF2α and eIF3, as well as the absence of density for eIF3b, NTD of eIF3c and CTD of eIF3h in the r48S AUG structure (Simonetti et al., 2020). We note that in contrast to our fully reconstituted h48S complexes, r48S AUG contains ABCE1, a ribosome recycling factor (Pisarev et al., 2010; Schuller and Green, 2017).

**Figure 3.**
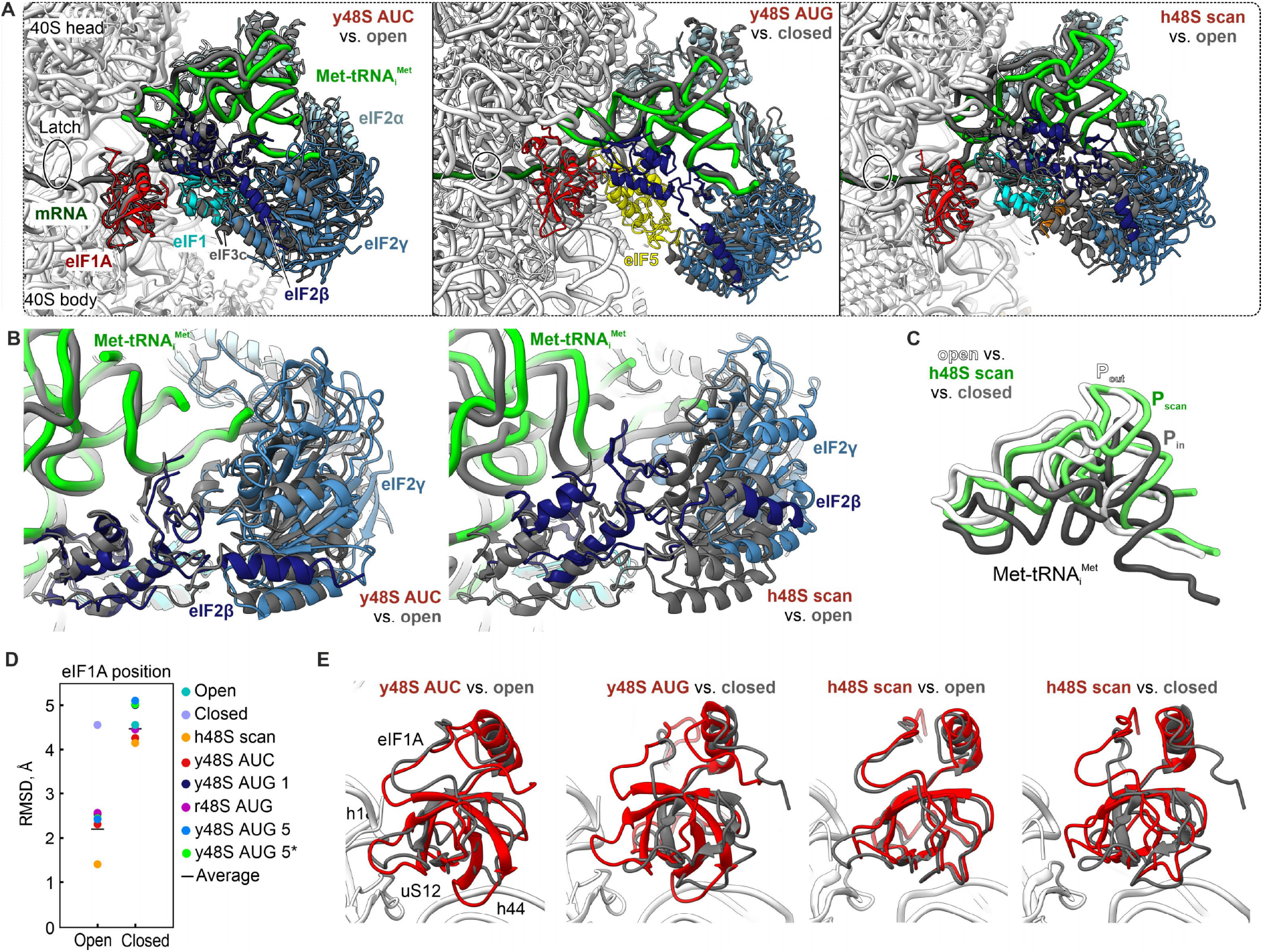
Structural dynamics of 48S IC. **(A)** Superposition of h48S AUG open and closed with other reported 48S IC structures. h48S AUG: IFs and tRNA are shown in dark grey; for visual clarity, the 40S subunit was removed after alignment. In structures used for comparison, IFs and tRNA are shown in color, 40S subunit in white. Structures used for comparison are y48S open with near-cognate AUC start codon (y48S AUC, PDB 3JAQ, (Llacer et al., 2015)), y48S closed with eIF5 bound (yellow) and cognate AUG start codon (y48S AUG, PDB 6FYX)) (Llacer et al., 2018), and h48S scan (PDB 6ZMW, (Brito Querido et al., 2020)). Note the stabilization of eIF2β in the decoding center of 48S ICs with eIF1 or eIF5 bound. eIF2β appears to dissociate from the decoding center in the closed h48S AUG IC, which has neither eIF1 nor eIF5 bound. **(B)** Comparison of eIF2β and tRNA body position and interactions in the present open state (dim grey) vs. reported 48S ICs (colored). **(C)** Position of initiator tRNA in the present open (P_out_) and closed (P_in_) states vs. the position in h48S scan (P_scan_). **(D)** Unique rearrangement of eIF1A in h48S AUG upon start codon recognition. Plot: Root-mean-square deviations (RMSDs) between the Cα-backbone of eIF1A in the present h48S AUG open and closed states and the reported 48S structures. y48S AUC, near-cognate yeast 48S IC with mRNA and eIF1 bound (PDB 3JAQ, (Llacer et al., 2015)); y48S AUG 1, cognate yeast 48S IC with cognate mRNA and eIF1 bound (PDB 3JAP, (Llacer et al., 2015)); y48S AUG 5, cognate yeast 48S IC with eIF5 bound in state C1 (PDB 6FYX, (Llacer et al., 2018); y48S AUG 5*, cognate yeast 48S IC with eIF5 bound in state C2 (PDB 6FYY, (Llacer et al., 2018)); r48S AUG, rabbit 48S IC with cognate β-globin mRNA (Simonetti et al., 2020); h48S scan, non-cognate human 48S IC (PDB 6ZMW, (Brito Querido et al., 2020)). **(E)** Superposition of eIF1A in present states (dim grey) vs. eIF1A in reported 48S ICs (red). Notably, the position of eIF1A is similar in all 48S ICs, except the present closed h48S AUG IC, which adopts a substantially different position.

Comparison between the structures of human 48S complexes reveals that h48S AUG open and closed described here differ substantially from the h48S·scanning complexes (Brito Querido et al., 2020). The orientation of the 40S head domain in h48S·scan (Figure 2 – figure supplement 1) renders partial opening of the decoding center and mRNA latch and a conformation of the TC distinct from open and closed states (Figures 2B, 3A and Figure 3 – figure supplement 1). Similar to the h48S AUG open, Met-tRNA_i_^Met^ in the scanning complex does not interact with the start codon (Figure 2B). However, due to the particular domain arrangement, the anticodon stem-loop of tRNA_i_^Met^ during scanning adopts a position half-way between P_out_ and P_in_ (P_scan_) enabling formation of a contact with eIF1 (Figure 3B,C, Figure 2D and Figure 3 – figure supplement 1). In comparison to the open state, the position of eIF1 in h48S scan is slightly shifted. In both h48S AUG open and h48 scan, eIF1 interacts with the N-terminal insertion of eIF3c (Figure 2 – figure supplement 1C). In contrast, the helix-turn-helix domain of eIF2β is found 10 Å away from the decoding center in the scanning complex compared to the open complex. The TC, in particular eIF2α and eIF2γ, move in the same direction as eIF2β and the contacts of eIF2β with tRNA_i_^Met^ are shifted from the anticodon-stem loop in the open state to the D loop region in the scanning complex (Figure 3A,B). In the closed state, eIF2α and eIF2γ move in a different direction than in h48S scan and towards the 40S body due to the simultaneous dissociation of eIF2β from the decoding center and tRNA_i_^Met^ docking onto the start codon (Figure 3 – figure supplement 1). The overall conformation of eIF3 is very similar in the human 48S complexes in the scanning, open and closed states, except for eIF3j subunit, which is visible only in the scanning complex.

A major difference between h48S AUG closed and all other reported 48S structures is the change in the position of eIF1A (Figure 3D,E), with a root-mean-square deviation (RMSD) of about 4.5 Å to any of these other structures. In the majority of structures, the position of eIF1A is similar to that in our open h48S AUG PIC conformation, with an RMSD of only ~2 Å. Considering the crucial role of eIF1A in start site selection (Fekete et al., 2007; Llacer et al., 2018; Maag et al., 2006), the unique position of eIF1A in our h48S AUG closed structure may represent an intermediate that forms after codon recognition, but prior to further steps on the path of 80S IC assembly.

### 48S IC conformations probed by time-resolved fluorescence measurements

Our findings showing the inherent structural heterogeneity of h48S AUG have prompted us to investigate the ribosome population distribution by time-resolved fluorescence methods. Following the previous work on y48S IC, we used eIF1A as a reporter for conformational changes in the complex (Llacer et al., 2015; Llacer et al., 2018; Maag et al., 2006; Maag et al., 2005). eIF1A is one of the first factors to bind to and one of the last to dissociate from the 40S subunit upon 80S IC formation (Acker et al., 2009; Fringer et al., 2007). Furthermore, the observed reorientation of eIF1A in h48S AUG closed (Figure 2F) is likely to change the local environment of eIF1A, which can be used to monitor the 40S subunit conformations using an environmentally sensitive fluorophore. To label different regions of eIF1A, we introduced single cysteine residues at positions N4, S74 or T120, which are located at the N-terminal tail (NTT), oligonucleotide/oligosaccharide-binding (OB) domain, and the C-terminal tail (CTT) of eIF1A, respectively (Figure 4A), and labeled these eIF1A variants with AlexaFluor555 (Alx555). All eIF1A derivatives are active in promoting 48S IC assembly and 80S EC formation as indicated by toe-printing assay and peptide bond formation with efficiency similar to that in the presence of WT eIF1A (Figure 1 – figure supplement 2D and Figure 4 – figure supplement 1).

**Figure 4.**
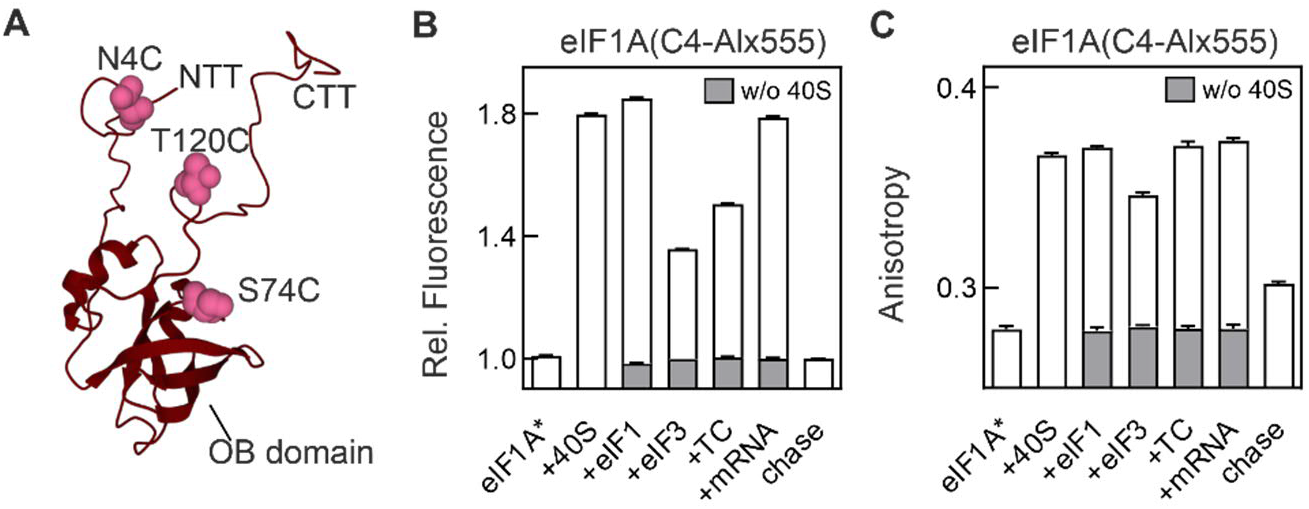
Monitoring the compositional and conformational dynamics of h48S using fluorescence-labeled eIF1A(C4-Alx555). **(A)** Structure of eIF1A (PDB 1D7Q; (Battiste et al., 2000)). Alexa 555-labeling positions (pink) are shown in the NTT, the OB domain, and the CTT of eIF1A. **(B), (C)** Relative fluorescence intensity **(B)** and anisotropy **(C)** changes of eIF1A(C4-Alx555). 48S IC was assembled by sequential addition of factors as indicated. In **(B)**, fluorescence intensity of free eIF1A(C4-Alexa555) is set to 1.0. Fluorescence intensity and anisotropy measured in the absence of the 40S subunit (w/o 40S) are shown in dark grey. Error bars represent standard deviations of 5 technical replicates (N=5).

We first tested whether the fluorescence intensity of labeled eIF1A is sensitive to compositional changes at different steps of 48S IC assembly. Each one of the labeled eIF1A derivatives shows a distinct fluorescence signature upon binding to the 40S subunit and subsequent stepwise addition of eIFs (Figure 4B and Figure 4 – figure supplement 2). The magnitude of the fluorescence change depends on the labeling position. For example, fluorescence of eIF1A(C4-Alx555) increases by 80% upon binding to the 40S subunit and is not further altered by eIF1 recruitment, but decreases considerably upon binding of eIF3 (Figure 4B). Addition of TC has only a small effect. Upon binding of mRNA with an AUG start codon, fluorescence intensity increases again, consistent with the expected rearrangement upon start codon recognition. Chase of eIF1A(C4-Alx555) with excess amount of unlabeled eIF1A restores the fluorescence value before binding to the 40S subunit, demonstrating that the signal results from a reversible interaction of eIF1A with the 40S subunit. In the absence of the 40S subunit, the fluorescence of eIF1A(C4-Alx555) does not change when other eIFs are added, showing that the observed fluorescence changes reflect the 48S IC assembly (Figure 4B). The fluorescence profiles of eIF1A(C74-Alx555) and eIF1A(C120-Alx555) are somewhat different from eIF1A(C4-Alx555), e.g., eIF1A(C74-Alx555) is particularly sensitive to eIF1 recruitment, whereas eIF1A(C120-A555) monitors 40S subunit and TC recruitment (Figure 4 – figure supplement 2). We further validated the fluorescence intensity approach by measuring anisotropy of eIF1A(C4-Alx555) (Figure 4C). The addition of 40S subunit to eIF1A(C4-A555) leads to an anisotropy increase due to formation of a stable 40S·eIF1A complex. Addition of eIFs and mRNA does not alter anisotropy, indicating that eIF1A remains bound to the 40S subunit throughout the 48S IC assembly. A small, but significant, anisotropy decrease with eIF3, together with the pronounced decrease of fluorescence intensity suggests that eIF3 binding to the 40S–eIF1A–eIF1 complex loosens eIF1A binding; this effect is reversed upon TC binding. Anisotropy decreases only when dissociation of eIF1A(C4-Alx555) is induced by the addition of excess unlabeled eIF1A (chase) or upon 80S formation induced by addition of eIF5B and the 60S subunit (see below), in agreement with the notion that eIF1A dissociates during 60S subunit joining (Acker et al., 2009; Fringer et al., 2007). In the following, we used the changes in eIF1A fluorescence intensity to follow structural and compositional rearrangements upon assembly of translation initiation complexes.

### h48S rearrangements on the pathway to start codon selection

Previous work on yeast initiation has shown that eIF1A dissociation kinetics is indicative of 40S subunit conformations (Llacer et al., 2018; Maag et al., 2006; Maag et al., 2005), which prompted us to use a similar approach to study how codon recognition affects the complex conformation in the mammalian system. We first measured the dissociation rates of fluorescence-labeled eIF1A from h48S CUC, h48S AUC and h48S AUG complexes upon mixing h48S with excess unlabeled eIF1A in a stopped-flow apparatus (Figure 5A and Table 1). To avoid potential bias due to the labeling position, we carried out these experiments with three eIF1A derivatives labeled at different sites of the protein, which showed very similar effects (Figure 5B-F and Table 1). Dissociation of eIF1A from the h43S PIC, h48S CUC and h48S AUC follows single exponential kinetics, with the rates of 0.01 s^-1^, 0.007 s^-1^, and 0.03 s^-1^, respectively. The dissociation is slow on a physiological scale, consistent with the notion that eIF1A remains bound to the 40S subunit until the 60S subunit joining (Acker et al., 2009; Fringer et al., 2007). In the simplest model, the single exponential kinetics suggests that the major contacts of eIF1A are similar on most of 40S subunits in the population, i.e. the h43S PIC and h48S PICs representing the scanning (h48S CUC) or partial codon reading (h48S AUC) comprise homogeneous populations with respect to the decoding center conformation. In contrast, chase of eIF1A from h48S AUG shows two-exponential dissociation kinetics (Figure 5E,F and Table 1), indicating the presence of two distinct populations of complexes. The slow dissociation (0.005 s^-1^) from h48S AUG is in the same range as from scanning h48S CUC. The additional kinetic phase has a much higher rate (0.11 s^-1^) and appears only in the presence of the cognate AUG codon, suggesting that start codon recognition induces rapid dissociation of eIF1A from a population of h48S IC. The presence of two distinct h48S AUG populations is consistent with the cryo-EM reconstructions presented above (Figure 1). It is therefore likely that the kinetic population with a rapid eIF1A dissociation that appears upon AUG codon recognition represents h48S AUG closed that is predominant in the cryo-EM sample and yields stable 48S IC in the toe-printing analysis (Figure 1 – figure supplement 2C), whereas the population that releases eIF1A more slowly corresponds to h48S AUG open population.

**Figure 5.**
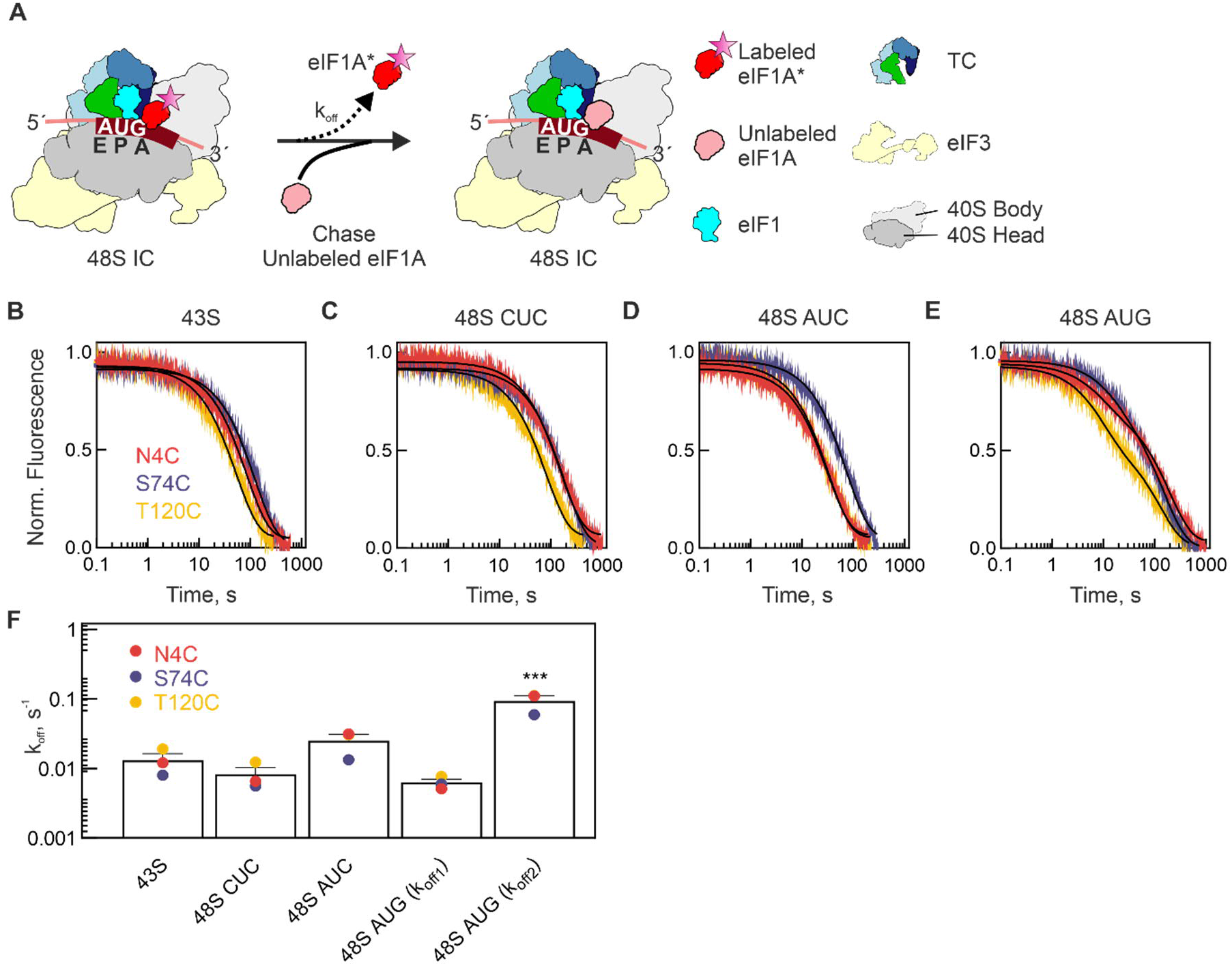
Dynamics of 48S IC upon start codon selection. **(A)** Schematic of the chase experiment. The 43S PIC or 48S IC (15 nM) assembled from 40S subunits with eIF1, TC, eIF3, eIF4A, eIF4B, fluorescence-labeled (*) eIF1A, with or without mRNA was rapidly mixed with excess of unlabeled eIF1A (1.8 μM) in the stopped-flow apparatus. **(B-E)** Time courses of eIF1A dissociation from 43S PIC **(B)**, 48S CUC **(C)**, 48S AUC **(D)** and 48S AUG IC **(E)** measured using eIF1A labeled at different positions. Black lines show results of one-exponential **(B-D)** or two-exponential **(E)** fitting. Fluorescence intensity change was normalized to 0-1 range. Time courses represent averages of 5 technical replicates (N=5). **(F)** Summary of the eIF1A dissociation rates from different initiation complexes (B-E). Bar graphs show average values from three different reporters with the standard deviation (N=3). For each reporter, the standard deviation of the measurement is smaller than the symbol size. ***indicates that the k_off_2 of 48S IC (AUG) measured with each eIF1A variant is significantly different from other k_off_ values for the same variant as compared using a two-tailed t-test (P-value <0.001). Note the logarithmic scale of the Y-axis.

**Table 1.**
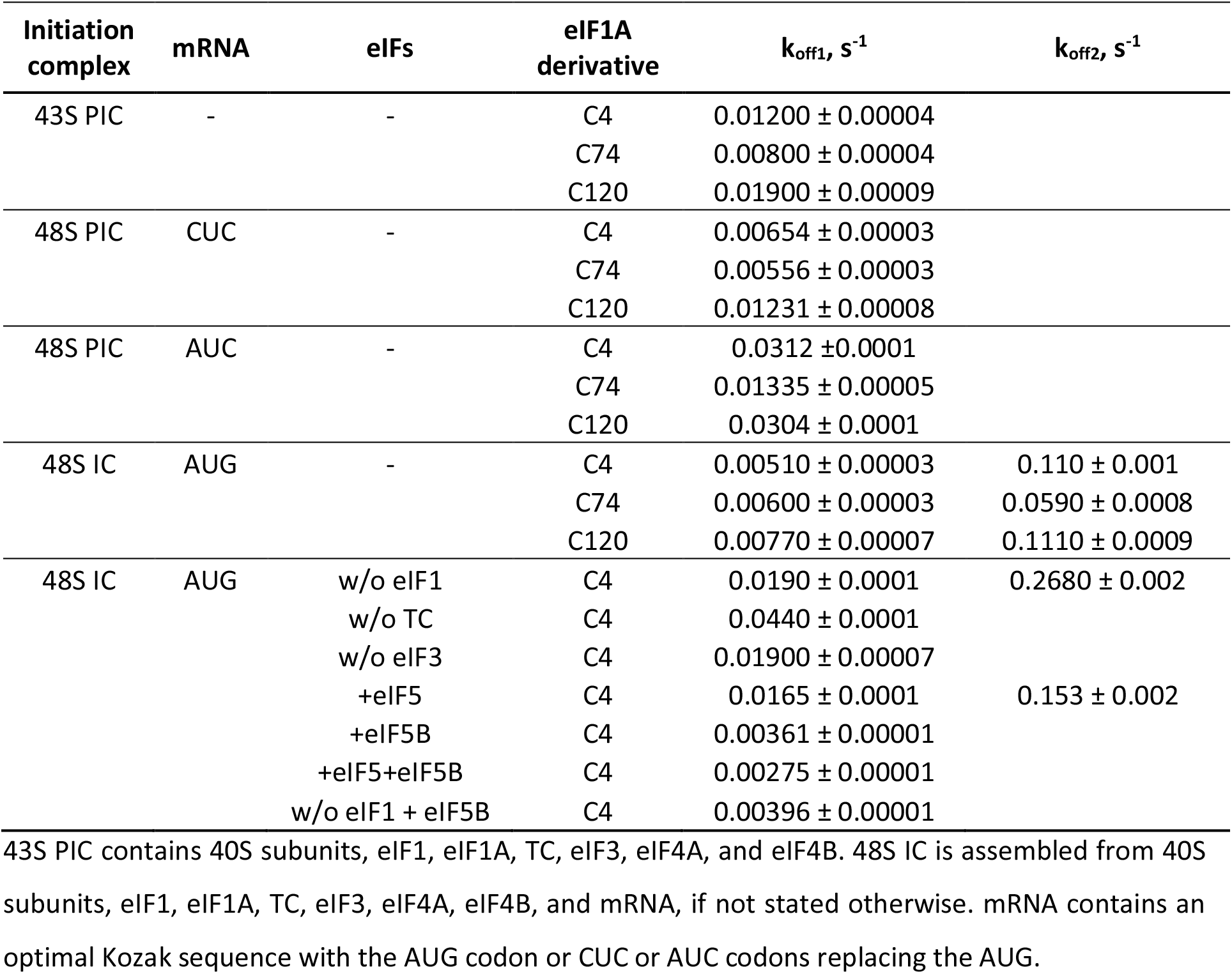
Summary of eIF1A dissociation rates from h43S PIC and various h48S complexes.

Our results suggest two interesting differences to the yeast system, where similar experiments were performed by measuring anisotropy of fluorescence-labeled eIF1A. First, eIF1A dissociation from y48S complex is biphasic irrespective of the initiation start codon. Second, AUG recognition results in a preferential stabilization of eIF1A on y48S IC (Llacer et al., 2018; Maag et al., 2006; Maag et al., 2005), suggesting that start codon recognition in yeast leads to tighter binding of eIF1A, rather than weaker binding which we find in the human system. In part, this may be due to the compositional differences between y48S IC and our h48S IC, as yeast complexes used in kinetic studies were mostly assembled in the absence of eIF3. This is possible, because eIF3 is dispensable for translation initiation on model mRNAs in yeast, but not feasible in mammalian system, where eIF3 is essential for h48S IC formation (Figure 1 – figure supplement 2B) (Majumdar et al., 2003; Pestova and Kolupaeva, 2002). In addition, y48S IC was assembled in the presence of eIF5, which is dispensable for h48S IC assembly (Figure 1 – figure supplement 2A,B)(Pestova and Kolupaeva, 2002). These differences prompted us to further investigate how the ribosome population distribution and eIF1A dissociation depend on the set of eIFs bound to h48S complexes.

### eIFs modulate ribosome population distribution

First, we followed eIF1A dissociation from h48S IC lacking one of the essential eIFs. In the absence of TC or eIF3, eIF1A release is relatively fast and the time courses are single exponential, indicating a uniform ribosome conformation with respect to eIF1A binding. In the mammalian system, TC and eIF3 are both required for start codon recognition, and in their absence, h48S PIC does not form a stable complex on the AUG codon, as indicated by the lack of the respective characteristic band in the toe-printing assay (Figure 1 – figure supplement 2B). TC stabilizes eIF1A binding on the 43S PIC in both yeast and mammals (Algire et al., 2002; Maag et al., 2005; Sokabe and Fraser, 2014), which explains why eIF1A dissociation is faster in the absence than in the presence of TC (Figure 6A,C and Table 1). eIF3 may affect eIF1A binding indirectly, e.g. by stabilizing eIF1 in the open state of the 40S subunit (Figure 3 – figure supplement 1A) (Majumdar et al., 2003; Sokabe and Fraser, 2014), which would also explain why the lack of these interactions results in a more rapid eIF1A dissociation.

**Figure 6.**
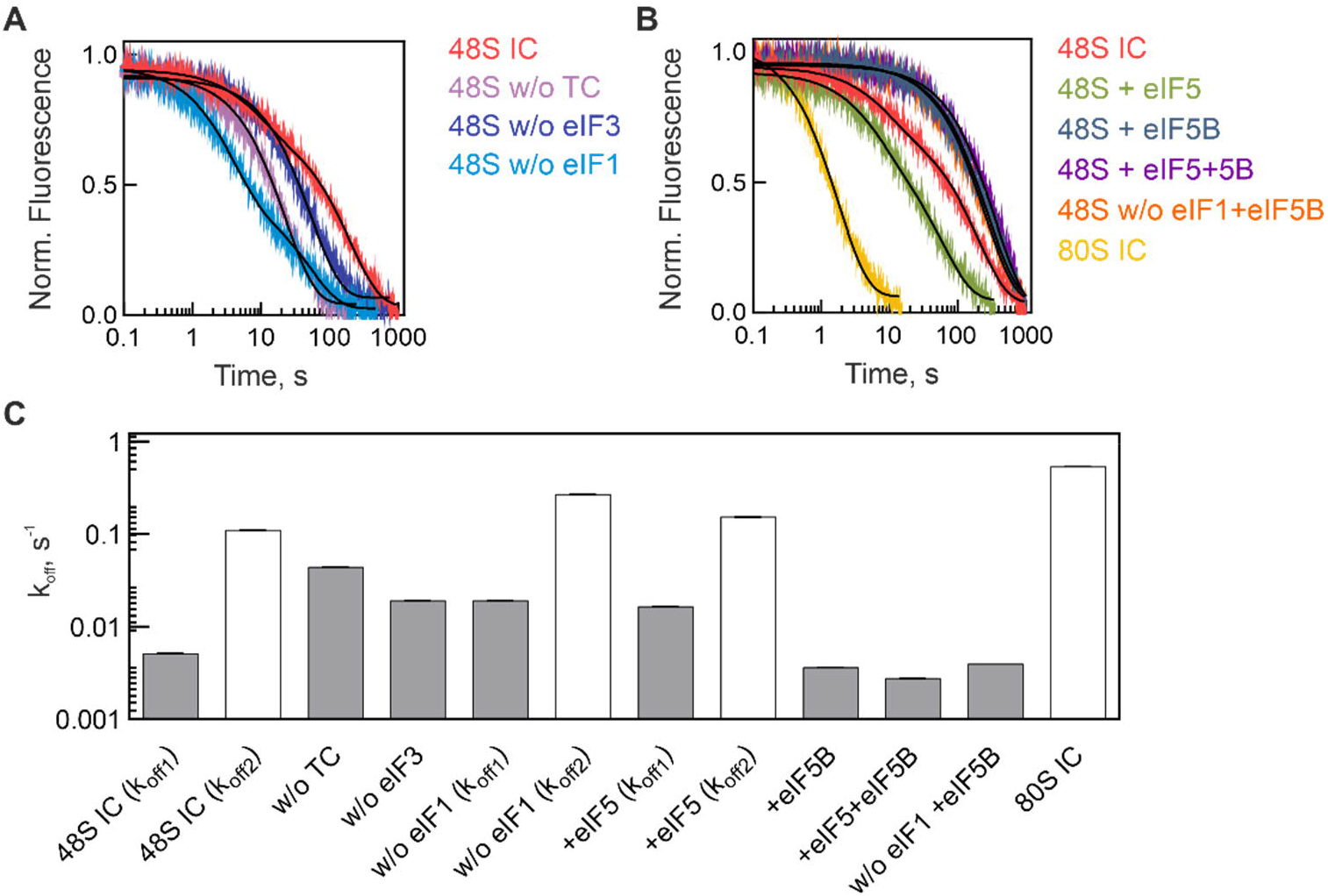
Dynamics of h48S IC. **(A)** Time-courses of eIF1A dissociation from the partial h48S IC lacking eIF1, TC or eIF3. Fitting of the time courses was single-exponential for complexes lacking TC or eIF3 and two-exponential for h48 IC with or without eIF1. Fluorescence change is normalized to 0-1 range. Time courses are averages of five technical replicates (N=5). **(B)** Time-courses of eIF1A dissociation from 48S IC in the presence of eIF5 or eIF5B or from 80S IC. Dissociation of eIF1A from the 80S complex was induced by mixing 60S subunits with the 48S AUG in the presence of eIF5 and eIF5B. Time courses in the presence of eIF5 were analyzed by two-exponential fitting; those with eIF5B, eIF5 and eIF5B, w/o eIF1+eIF5B, and 80S IC were analyzed by one-exponential fitting. Fluorescence change is normalized to 0-1 range. Time courses are averages of five technical replicates (N=5). **(C)** Summary of the k_off_ values. Rates of single-exponential reactions as well as slow phases of two-exponential time courses are shown in grey, the fast phases and dissociation from 80S in white bars. Error bars (very small) show standard deviation of five technical replicates (N=5).

In contrast to other factor omission experiments, eIF1A dissociation from h48S AUG lacking eIF1 is biphasic and both dissociation rates are somewhat faster than from the complete h48S IC (Figure 6A,C and Table 1). The increase in the dissociation rates is expected, because eIF1 and eIF1A are known to stabilize each other’s binding to the 40S subunit (Maag and Lorsch, 2003; Majumdar et al., 2003; Pestova et al., 1998; Sokabe and Fraser, 2014). Notably, we found a stable 48S IC toe-print on the AUG codon in the absence of eIF1 (Figure 1 – figure supplement 2D). This is consistent with the notion that eIF1A can compensate for the lack of eIF1 during 48S IC assembly on a model mRNA with an optimal initiation start site, as used in this study (Fekete et al., 2007; Pestova and Kolupaeva, 2002). This also explains why h48S IC lacking eIF1 can recognize the AUG codon, leading to formation of the characteristic two-population distribution of the 40S subunits in the complex (Figure 6A,C and Table 1).

Next, we studied the effect of eIF5 and eIF5B (Figure 6B,C and Table 1). eIF1A dissociation from h48S AUG in the presence of eIF5 is biphasic and the dissociation rates are comparable to those from h48S IC without eIF1 (Fig 6A,C and Table 1). The latter is in agreement with the finding that eIF5 displaces eIF1 upon start codon recognition (Llacer et al., 2018), but the observed destabilization of eIF1A kinetics on h48S AUG is at variance to the yeast system, where eIF1A binds more tightly upon eIF5 addition (Llacer et al., 2018; Maag et al., 2006; Maag et al., 2005). Finally, addition of eIF5B stabilizes the h48S AUG in a single major population with tight eIF1A binding (Figure 6B,C and Table 1). eIF5B compensates the destabilizing effect of eIF5 and reverses the eIF1 omission effect, suggesting that eIF5B acts after the remodeling of the decoding center by eIF5. The h48S IC with eIF5B bound is the last intermediate on the pathway of translation initiation before the 60S subunit docking, which remodels the complex to an 80S IC ready to start translation. The 60S subunit joining triggers fast release of eIF1A from the 80S IC with a rate of 0.53 s^-1^. Overall, these results support the notion that the two-population distribution of h48S AUG is characteristic for complexes upon start codon recognition in the mammalian system.

## Discussion

The results of our experiments suggest how start codon recognition modulates the structure of human h48S IC (Figure 7). In h43S PIC, which we characterize by kinetic experiments using eIF1A as reporter, 40S subunit is found in a single predominant conformation that binds eIF1A tightly. This characteristic state of the complex is maintained throughout mRNA scanning. Partial recognition of the near-cognate codon AUC by Met-tRNA_i_^Met^ has a small destabilizing effect on eIF1A binding. Start codon recognition changes the conformational distribution in the ensemble, and both kinetic and structural studies consistently show two conformations of h48S AUG. The structure of the open h48S AUG PIC shows a small fraction of complexes that do not undergo codon-anticodon interaction and bind eIF1A tightly. On the majority of ribosomes, however, start codon recognition induces structural remodeling of the decoding site that induces 40S domain closure, movement of Met-tRNA_i_^Met^ into the P_in_ conformation, displacement of eIF1 and eIF2β, as well as a rearrangement of eIF1A that results in faster dissociation rate of eIF1A from the ribosome. Binding of eIF5 has an additional, albeit small, destabilizing effect, but the resulting complexes still comprise two kinetic populations. Recruitment of eIF5B, which is a prerequisite for the efficient 60S subunit docking, induces a rearrangement to a predominant 40S conformation to which eIF1A binds tightly, potentially owing to a direct contact between eIF5B and eIF1A (Acker et al., 2006; Acker et al., 2009; Lin et al., 2018). Finally, eIF1A dissociates rapidly from the 80S IC, probably following GTP hydrolysis by eIF5B (Acker et al., 2009; Lee et al., 2002; Pestova and Kolupaeva, 2002; Pestova et al., 2000) after 60S joining (Figure 7).

**Figure 7.**
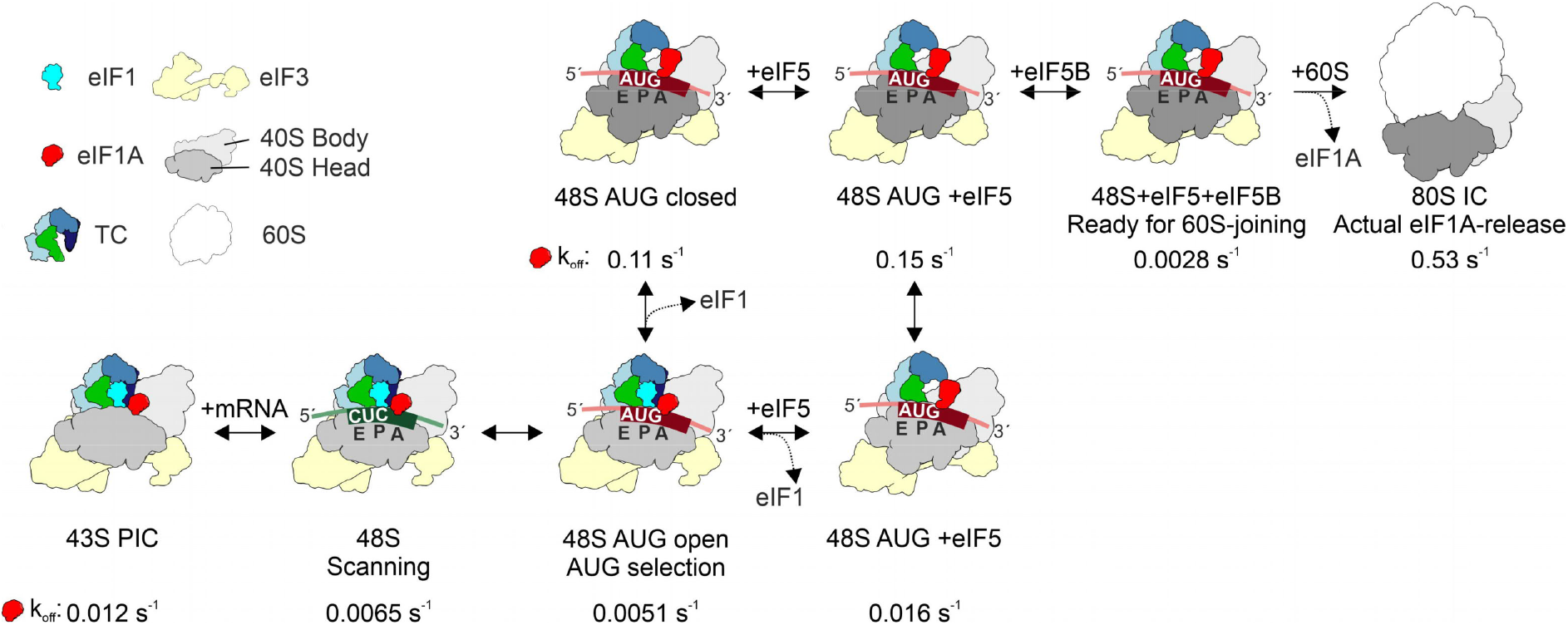
Kinetic model of late-stage human 48S IC assembly. 43S PIC and 48S scanning complex comprise uniform populations as monitored by eIF1A dissociation kinetics; eIF1A dissociation rates (k_off_) are indicated. In contrast, h48S AUG complexes comprise two populations that correspond to h48S AUG open (prior to codon recognition, with low k_off_ for eIF1A, labeled with light 40S head) and h48S AUG closed (after codon recognition, with high k_off_ for eIF1A, labeled with dark 40S head). Upon codon recognition, eIF1 is displaced, which can occur spontaneously (as indicated by the cryo-EM structures in this paper) or facilitated by eIF5. In the mammalian system, these rearrangements depend on eIF3. A partial codon recognition complex akin to that formed in y48S AUC has a low eIF1A dissociation rate (0.03 s^-1^) and may transiently form also upon AUG recognition, but is not captured by structural studies and therefore not indicated in the scheme. Binding of eIF5B stabilizes a single population which binds eIF1A tightly. Finally, upon 60S joining, eIF1A rapidly dissociates from 80S IC. The binding sites of eIF5 and eIF5B are not shown for visual clarity. The k_off_ values in the schematic are derived from measurements with eIF1A(C4-Alx555).

The structure of h48S AUG open differs from the scanning conformation of h48S (Brito Querido et al., 2020), suggesting that the two states are functionally distinct or at least represent different snapshots at the scanning pathway. In both structures, the codon-anticodon interaction is not established, but the 40S head domain is tilted to a different degree, resulting in a shift of Met-tRNA_i_^Met^ position. On the other hand, open h48S AUG PIC is similar to y48S AUC, except for the partial codon-anticodon interaction in the near-cognate yeast complex, which is not found in h48S AUG open. Because h48S AUG open contains a full complement of eIFs required to form h48S IC, this intermediate is unlikely to be an inactive off-pathway complex. With these comparisons in mind, we suggest that h48S AUG open represents ribosomes that attempt to read the start codon, but before the tRNA succeeds to form the first base-pair of the stable codon-anticodon complex. The observed 40S domain opening compared to the scanning conformation may facilitate the accommodation of Met-tRNA_i_^Met^ and mRNA in the decoding center. Thus, if partial AUC codon recognition complexes are similar in yeast and human systems, a likely order of complexes on the translation initiation pathway is 43S PIC –> scanning h48S CUC PIC –>h48S AUG open PIC –> partial codon recognition 48S AUC PIC –> closed h48S AUG IC (Figure 7). We note, however, that there are kinetic differences between the yeast and human 43S and 48S PICs, as revealed by the eIF1A dissociation assay. While h43S PIC, as well as the scanning and partial codon recognition h48S PIC comprise a single major kinetic population, all yeast complexes entail two populations (Llacer et al., 2018; Maag et al., 2006; Maag et al., 2005). The structural basis for the two distinct conformations in y48S PIC is not clear, as structural studies reveal a single major ribosome population of y48S AUC PIC or y48S AUG IC, possibly due to the use of the tRNA_i_^Met^ mutant that stabilized the P_in_ state (Llacer et al., 2015).

The population of the ribosomes that have formed stable codon-anticodon interactions adopted a closed conformation. They are structurally similar to other reported 48S IC complexes from yeast and mammals (Eliseev et al., 2018; Llacer et al., 2015; Llacer et al., 2018; Simonetti et al., 2020). Surprisingly, the presence of eIF1A in r48S IC appears to depend on the type of the mRNA, e.g., the complex with the histone H4 mRNA displays no eIF1A density, whereas the complex with β-globin mRNA shows eIF1A in a state similar to that in our open h48S AUG state (Simonetti et al., 2020). Several other structures of mammalian complexes lacking one or more of the essential factors have been reported, but are not discussed here (Eliseev et al., 2018; Simonetti et al., 2016) (Simonetti et al., 2020).

The destabilizing effect of start codon recognition on eIF1A binding to h48S AUG IC is at odds with the previous results obtained in the yeast system, where codon-recognition was shown to increase the binding stability of eIF1A. One potential explanation would be the presence of eIF5 in yeast complexes, as we initially did not use eIF5 to assemble h48S AUG IC. We note that potential eIF5-dependent effect would be due to eIF5 binding, rather than GTP hydrolysis, as kinetic experiments in the yeast system were carried out in the presence of a non-hydrolysable GTP analog (Llacer et al., 2018; Maag et al., 2006). Our complexes contain GTP, which is hydrolyzed only upon addition of eIF5 (Majumdar and Maitra, 2005; Unbehaun et al., 2004); thus, in both h48S AUG and y48S AUG, eIF2 is expected to be in the GTP-bound form. Our kinetic measurements suggest that in the human system addition of eIF5 (or omitting eIF1, which should mimic the remodeling effect of eIF5 on the decoding center) did not stabilize eIF1A binding on either of the two ribosome populations. These findings suggest that the difference in kinetic stability of eIF1A in the mammalian vs. yeast system is not due to the presence or absence of eIF5, but must depend on other factors, e.g. the presence of eIF3 or the details of interactions that are difficult to discern at the current resolution of existing cryo-EM structures. In mammals, eIF1 stabilizes the binding of eIF1A and inhibits premature GTP hydrolysis by eIF2. AUG recognition relieves this inhibition due to replacement of eIF1 by eIF5 and thereby destabilizes eIF1A binding. Thus, the opposite effects of codon recognition on the stability of eIF1A binding in mammals and yeast may be related to the different ways by which eIF1 controls the GTPase activity of eIF2 in these organisms.

Another difference to the yeast system concerns the effect of eIF3. In yeast, the eIF1A binding is independent of the presence of eIF3 (Maag et al., 2006). In contrast, in the mammalian system eIF3 is essential for h48S IC formation (Kumar et al., 2016; Pestova and Kolupaeva, 2002; Sokabe and Fraser, 2014; Sokabe et al., 2012). eIF3 together with eIF1 stabilize eIF1A binding to the 40S subunit and are essential for recruiting the TC to the 43S PIC and for scanning (Kumar et al., 2016; Majumdar et al., 2003; Pestova and Kolupaeva, 2002; Sokabe and Fraser, 2014). eIF3 contacts the mRNA 5’ UTR upstream of the start codon and protects 17 nt of the mRNA in the vicinity of the mRNA exit channel in h48S IC (Pisarev et al., 2008). Interactions of eIF3c with eIF1, and at the later initiation steps with eIF5, are important for start site selection (Obayashi et al., 2017; Valasek et al., 2004). Because eIF1 stabilizes eIF1A binding and the eukaryotic-specific α-helical element of eIF3c interacts with eIF1 (Figure 2D), this interaction network may stabilize eIF1A binding, consistent with the low dissociation rate of eIF1A from h48S AUG open. Furthermore, the α-helical element of eIF3c bridges between its CTD in the eIF3 core and its NTD on the 40S intersubunit interface, where it blocks 60S subunit joining (Figure 2 – figure supplement 1C). In h48S AUG closed, the density of eIF3c α-helical element is not observed, indicating a structural rearrangement of the eIF3 subunits upon start codon recognition, which might affect eIF1A binding indirectly through release of eIF1-eIF3c contacts. These unique features of mammalian eIF3 may explain its specific role in modulating the mammalian 48S IC conformation compared to y48S IC.

In summary, the principle conformational rearrangements of the 48S PIC induced by start codon recognition, e.g., 40S subunit closure, movement of Met-tRNA_i_^Met^ and the displacement of eIF1 and eIF2β from the decoding site, appear to be conserved between lower and higher eukaryotes; however, their timing and regulation are notably different. Upon codon recognition, eIF1A binding is destabilized in human, but stabilized in yeast system. eIF3 is essential for start codon recognition in the mammalian system, but not in yeast. eIF5 and GTP hydrolysis by eIF2 are not necessary for codon recognition in mammalian system, consistent with the notion that eIF5 is not essential for reconstitution of h48S IC. In mammals, start codon recognition relieves the eIF1-gated inhibition of GTP hydrolysis by eIF2 in the presence of eIF5, which is necessary to remove eIF2 from the complex (Majumdar and Maitra, 2005; Unbehaun et al., 2004). In contrast, in yeast eIF5 plays a key role not only in promoting GTP hydrolysis and eIF2 dissociation, but also in triggering Pi release from eIF2 upon start codon recognition (Algire et al., 2005; Jennings and Pavitt, 2010; Nanda et al., 2013; Saini et al., 2014). In human system, eIF1A binding is stabilized by eIF5B after codon recognition, whereas in yeast eIF5B binding appears to have no further effect on eIF1A (Acker et al., 2009). The functional consequences of these differences in timing and regulation of start codon selection are not known, but may be related to the essential role of eIF3 in translation regulation in mammals (Gomes-Duarte et al., 2018; Lee et al., 2015; Lee et al., 2016; Zeman et al., 2019), linked to complex networks of interactions with auxiliary regulators that play important roles in health and disease in mammalian cells.

## Materials and Methods

### Biochemical methods

#### Model mRNAs

The model mRNA (GGG CAA CAA CAA CAA GCU AGC CACAA CAA CAA CAA CAA CAA CAA CAA GUC GAC CAA CAA CAA CAA CAA CAA CAA CAA CAA CUC GAG CAA CAA CAA CAA CAA CAA CAA CAA CAA GGA UCC AAA ACA GAC CACC **<u>AUG/AUC/CUC</u>** GUA CGU UUC AAG GCU UGA GCC CUC GUC ACU GCC CUG UGG GGC AAG GUG ACU CUG GAA GAA GUU GGU GGU GAG GCC CUG GGC AGG CUG CAG AGU GUG AGG GAA GGU CUG GUU GUC UAC CAA) is constructed using pUC19 and amplified with forward (5’-ATCTAGAATTCTAATACGACTCACTATAGG-3’) and reverse (5’-TTGGTAGACAACCAGACCTTCC-3’) primers. The PCR product was transcribed *in vitro* using T7 RNA-polymerase and purified by a Hitrap Q HP column (GE Healthcare) with ethanol precipitation.

#### Met-tRNA_i_^Met^ preparations

Full-length human tRNA_i_^Met^ sequence is inserted into the pIDTSMART-AMP plasmid (Integrated DNA Technologies) and amplified with forward (5’-TAATACGACTCACTATAAGCAGAGTGG-3’) and reverse (5’-TGGTAGCAGAGGATGGTTTCGATC-3’) primers. tRNA_i_^Met^ is prepared by *in-vitro* transcription using T7 RNA-polymerase and purified as described above for mRNA. tRNA was aminoacylated in the presence of S100 extract from *E. coli* (in-house made (Milon et al., 2007)) with [^3^H]-methionine. [^3^H]Met-tRNA_i_^Met^ is purified by reverse-phase chromatography (Milon et al., 2007).

#### Purification of recombinantly-expressed eIFs (eIF1, eIF1A, eIF4A, eIF4B, and eIF5)

Sequences encoding full-length human eIF1, eIF1A, eIF4A, eIF4B, and eIF5 were cloned into the pET24a vector without a tag or into ptxB1 vector (eIF5) with a cleavable intein-linked His-tag. eIF1, eIF1A and eIF5 were expressed in *E. coli* BL21 codon-plus RILP (Agilent Technologies); eIF4A and eIF4B were expressed in *E. coli* Rosetta strain. Recombinant proteins were purified on a HiTrap SP HP column (GE Healthcare) and subsequently on a HiTrap Q HP column (GE Healthcare) using a 50–1000 mM KCl gradient in buffer (20 mM Tris-HCl, pH 7.5, 0.1 mM EDTA, 5% glycerol, 2 mM DTT) at 4 °C. For eIF4A, an additional HiTrap Blue column (GE Healthcare) was introduced after the HiTrap Q HP column. For eIF5, an additional Protino Ni-IDA column (Macherey-Nagel) was applied before the Hitrap SP HP column and the His-tag was cleaved with 50 mM DTT in buffer (20 mM Tris-HCl, pH 7.5, 0.1 mM EDTA, 5% glycerol, 100 mM KCl). Recombinant proteins were identified by SDS-PAGE (12%) and confirmed by mass spectrometry.

eIF1A derivatives (N4C, S74C, T120C) were prepared in the same way as described above. Purified eIF1A derivatives were labeled at 4 °C with 4-fold excess of Alexa Fluor™ 555 C2 maleimide (Alexa555, Thermo Scientific) following the manufacturer’s manual. To remove the excess free dye, eIF1A preparations were re-purified on a HiTrap SP HP column and the fractions containing labeled eIF1A visualized on a 15 % SDS-PAGE using a fluorescence scanner (Typhoon™ FLA7000 (GE Healthcare)).

#### Purification of native human factors (eIF2, eIF3, eIF5B, eEF1A and eEF2), 40S and 60S subunits

Native human initiation factors and ribosomes were prepared from HeLa cytoplasmic lysate as described (Pisarev et al., 2007). In brief, the polysomes were isolated from the HeLa cytoplasmic lysate pelleting through a 2 M sucrose cushion in buffer (20 mM Tris-HCl, pH 7.5, 20 mM KCl, 4 mM MgCl2, 2 mM DTT) by 18-hour centrifugation with a Ti 50.2 rotor at 45000 r.p.m. at 4 °C. After resuspension of the polysomes, ribosome-bound factors were washed-off by increasing KCl concentration to 0.5 M. Factor-free polysomes were pelleted by centrifugation in a Ti 50.2 rotor at 45000 r.p.m at 4 °C for 4.5 hr. Mixtures of factors were then precipitated by increasing ammonium sulfate concentration to 40% and after pelleting to 50%. The 40% ammonium sulfate fraction is used for eIF3 purification and the 40–50% ammonium sulfate fraction is a source for eIF2 and eIF5B. After ammonium sulfate precipitation, proteins were resuspended and factors were purified on a HiTrap SP HP column using a 100–1000 mM KCl gradient in buffer (20 mM Tris-HCl, pH 7.5, 0.1 mM EDTA pH 8, 5% glycerol, 2 mM DTT), followed by an additional purification step on a MonoQ 5/50 GL column (GE Healthcare) at 4 °C. The elongation factors, eEF1A and eEF2, were purified in a similar way from 30–70% ammonium sulfate fraction as described previously (Pestova and Hellen, 2005). The purity of native initiation factors was confirmed by gradient SDS-PAGE gels and mass spectrometry. Notably, purified native human eIF3 contains significant amount of eIF4G, which prompted us to supply additional eIF4A and eIF4B in the reconstituted complexes. This increased the initiation efficiency only slightly; nevertheless, we kept the factors in the reaction mixture for better comparison with other experiments.

For human ribosome purification, factor-free polysome fraction was resuspended in buffer (20 mM Tris-HCl, pH 7.5, 4 mM MgCl2, 50 mM KCl, 2 mM DTT) at 4 °C and treated with 1 mM puromycin to release nascent peptides at 37 °C for 10 min. 80S ribosomes were split into 40S and 60S subunits by increasing KCl concentration to 0.5 M at 4 °C. Mixtures of 40S and 60S subunits were applied to an ice-cold 10–30% sucrose gradient and separated by overnight centrifugation using an SW32 Ti rotor at 4 °C. Fractions containing 40S and 60S subunits were identified by 260 nm absorbance and confirmed by mass spectrometry.

#### 48S IC preparation

Human 48S complexes were assembled in buffer (20 mM Hepes, pH 7.5, 95 mM KOAc, 3.75 mM Mg(OAc)_2_, 1 mM ATP, 0.5 mM GTP, 0.25 mM spermidine, 2 mM DTT, 0.4 U/μL RiboLock RNase inhibitor) with 0.36 μM 40S subunits, 1.1 μM eIF1, eIF1A, eIF4A, and mRNA each, 0.54 μM eIF3 and eIF4B each, 0.72 μM eIF2, and 0.8 μM Met-tRNA_i_^Met^ at 37 °C for 10 min. To assemble 80S IC, additional 3.6 μM eIF5, 1.1 μM eIF5B and 0.72 μM 60S subunits were added after 48S IC formation at 37 °C for 10 min.

#### Toe-printing assay

In the toe-printing assay, 48S IC and 80S IC were assembled as described above, except for the mRNA concentrations, which was 0.2 μM. Toe-printing primer (Atto647N-GACCTTCCCTCACACTCTG) (0.05 μM) was added into the 48S IC and 80S IC mixtures in reverse-transcription buffer (0.5 mM dNTPs, 8 mM MgCl2, 0.15 U/μL SuperScript III reverse transcriptase (Invitrogen)) and the reaction was incubated for 45 min at 37 °C. The cDNA products were extracted with phenol:chloroform:isoamyl alcohol and analyzed on an 8% urea-PAGE using fluorescence scanner (Typhoon™ FLA7000, GE Healthcare).

#### Spectrofluorometer assays

To monitor fluorescence changes of eIF1A upon 48S IC formation, 0.06 μM labeled eIF1A variants were used in each measurement and components to assemble the 48S IC were added step-wise into the reaction mixture at the same concentration and condition as described above. 3.6 μM unlabeled eIF1A was added to chase the labeled eIF1A variants from 48S IC. After adding each component, the fluorescence intensity and anisotropy were measured at 25 °C with excitation at 555 nm and emission at 568 nm. 5 technical replicates were taken for each measurement. For determinations of K_d_ and K_i_ of eIF1A to the 40S subunit, 5 nM eIF1A(C4-Alx555) or 5 nM 40S–eIF1A(C4-Alx555) complex were mixed with increasing concentrations of the 40S subunit or unlabeled eIF1A, respectively. The anisotropy was measured in the same condition as described above.

#### Chase experiments

43S and 48S complexes were assembled as described above using 0.06 μM Alexa555-labeled eIF1A variants. Additionally, 0.05% bovine serum albumin was included in the buffer to prevent non-specific binding of labeled eIF1A to the cuvette walls. In the conditions of 48S +eIF5 or +eIF5B, additional 3.6 μM eIF5 or 1.1 μM eIF5B were included. The 48S complexes were rapidly mixed with 7.2 μM of unlabeled WT eIF1A at 25°C in the stopped-flow apparatus (SX-20MV (Applied Photophysics)). For dissociation of eIF1A from 80S IC, the 48S+eIF5+eIF5B complex was rapidly mixed with 5.4 μM 60S subunit at 25°C in a stopped-flow apparatus. The fluorescence intensity was recorded with 4000 time points in logarithmic spacing using 535 nm excitation and a 570 nm emission filter. 5 technical replicates of time courses were collected for each experiment, averaged, and analyzed by single or double exponential fitting by GraphPad Prism.

### Cryo-EM methods

#### GraFix and Cryo-EM grid preparation

Complex preparation for cryo-EM was carried out in buffer (20 mM Hepes, pH 7.5, 95 mM KOAc, 3.75 mM Mg(OAc)_2_, 1 mM ATP, 0.5 mM GTP) at 4°C. The complexes were stabilized for cryo-EM grid preparation by the GraFix approach (Kastner et al., 2008) with some modifications. Specifically, complexes were stabilized before gradient centrifugation by 30 min incubation with 2 mM of a mild crosslinking reagent bis(sulfosuccinimidyl)suberate (BS3, Sigma Aldrich) at room temperature. Subsequently, complexes were crosslinked upon ultra-centrifugation on a linear 10-40% sucrose gradient (total volume 4.4 ml run for 16 h at 138,000 x g) combined with linear gradients of 0-0.1% glutaraldehyde (EM grade 25%, Science Services GmbH, Munich, Germany) and 0-1.0 mM p-maleimidophenyl isocyanate (PMPI, ThermoFischer Scientific). PMPI was introduced because of its heterobifunctional activity in crosslinking RNA and proteins. The gradient was fractionated into 200 μl fractions and the crosslinking reaction was quenched using 100 mM aspartate (Sigma Aldrich) at pH 7.5. Sucrose was removed using Zeba Spin columns (ThermoFischer Scientific), which were pre-incubated with 0.1 ml/mg gelatin (Sigma Aldrich) and then washed with buffer to improve sample recovery. Cryo-EM grids were prepared by floating home-made continuous carbon on 40 μl sample in the wells of teflon block (custom-made). The sample-covered carbon was then adsorbed to an EM grid (Quantifoil R3.5/1, Jena Bioscience) and blotted for 9 s using a Vitrobot Mark IV (ThermoFisher, Eindhoven) operated at 4°C and 100% humidity.

#### Data acquisition

Cryo-EM data acquisition was performed using a Falcon III direct electron detector (ThermoFisher, Eindhoven) on a Titan Krios G1 microscope with 300 kV acceleration voltage equipped with an XFEG electron source and a Cs-corrector (CEOS, Heidelberg) aligned with the CETCORPLUS 4.6.9 (CEOS, Heidelberg) software package. The total dataset of 15,544 cryo-EM movies (4096 x 4096 pixels) with 20 frames each was acquired using EPU 1.11 (ThermoFisher Eindhoven) in integration mode and 1 s exposure time at total electron dose of 48 electrons per Å^2^ and a defocus range of 1.5-4.0 μm.

#### Data processing

Beam-induced motion correction with dose-weighting was performed with MotionCor2 (Zheng et al., 2017) using 5×5 patches. CTF parameters of the motion-corrected micrographs were determined by Gctf (Zhang, 2016) and 990,486 particles particle were selected with Gautomatch (K. Zhang, MRC-LMB, Cambridge). All subsequent image processing was performed using RELION 3.0. The selected particles were extracted at 2.32 Å/pixel and sorted in a hierarchical manner using RELION 3.0 (Figure 1 – figure supplement 3A). 2D classification reduced the total number of particles to 821,651, which were then refined and sorted for 40S head conformation by focused 3D classification with signal subtraction without alignment using a mask for the 40S head (top panel in Figure 1 – figure supplement 3B) revealing two major populations with an open and a closed conformation of the 40S head, respectively. Subsequently, each of the particle populations was sorted separately focusing on the region of the decoding center and ternary complex area by 3D classification with signal subtraction (bottom panel in Figure 1 – figure supplement 3B). The resulting particle populations were each re-extracted at the final pixel size of 1.16 A/pix, their CTF parameters were locally determined using CTF refinement and the particles were refined to high-resolution following the gold-standard procedure using soft solvent masks (Figure 1 – figure supplement 3C). Global amplitude sharpening of the two final maps was performed using Phenix 1.16-3549 (Liebschner et al., 2019). To estimate the occupancy with eIF3, the final particle populations of the h48S open and closed state were each refined against the isolated 40S subunit to avoid any reference bias for eIF3 and then sorted into ten classes by focused classification with signal subtraction using a mask on the core of eIF3. In each case, all classes showed clear density for the eIF3 core indicating a high occupancy above 80% in both states.

#### Atomic model refinement

Initial models were obtained by rigid-body fitting individual chains of the structure from the human 48S complex in the scanning state (PDB 6ZMW, (Brito Querido et al., 2020)) into each of the two final cryo-EM maps using ChimeraX 1.2 (Pettersen et al., 2021), and subsequent manual adjustments and refinement in Coot 0.9.3 (Brown et al., 2015). Local atomic model refinement was done using low pass filtered to corresponding local resolutions of the maps (Figure 1 – figure supplement 3E) in Coot. For the eIF3 core atomic model refinement was done at 6 Å, and at 12Å for peripheral subunits, eIF2γ was fitted as rigid-body and refined with all-atom restraints at 12 Å, the 40S subunit, and the decoding center area of ternary complex was refined at the respective final resolutions, i.e. 3.7 Å and 4.7 Å, and eIF1A was refined at 6 Å resolution. The mRNA in both states was first modeled manually in Coot and then refined in ISOLDE 1.1.0 (Croll, 2018). Nucleotide bases for the mRNA in the open state were omitted, due to the weak density for the mRNA in this state. The resulting initial models were then automatically refined using phenix.real_space_refine (Liebschner et al., 2019) with secondary structure restraints for 10 macrocycles 500 iterations each.

### Crosslinking-mass spectrometry

The reconstituted 48S AUG IC was treated with either 2 mM BS3 (ThermoFisherScientific) or 2 mM LC-SDA (ThermoFisherScientific) for 1 h at RT. For LC-SDA crosslinking, the complex was dialyzed against buffer (20 mM Hepes, pH 7.5, 95 mM KOAc, 3.75 mM Mg(OAc)_2_, 1 mM ATP, 0.5 mM GTP) via a membrane filter (MF Membrane Filters, 0.025 μm VSWP, Merck) prior to crosslinking. LC-SDA crosslinked samples were irradiated with UV light (365 nm) for 5 min at 4°C. Crosslinking reaction with Bs3 or LC-SDA were quenched with 50 mM Tris-HCl pH 7.5 for 15 min. (Crosslinked) proteins were reduced with 10 mM dithiothreitol and subsequently alkylated with 40 mM iodoacetamide under standard conditions. Proteins were digested with the endoproteinase trypsin in an enzyme-to-protein ratio of 1:50 in the presence of 1 M urea at 37°C overnight. The reaction was terminated with 0.5% trifluoroacetic acid (TFA) (v/v), the peptide mixtures were desalted on MicroSpin Columns (Harvard Apparatus) following manufacturer’s instructions and vacuum-dried in a SpeedVac. Peptides were dissolved in 50 μL 30% acetonitrile (v/v) in water /0.1% (v/v) trifluoroacetic acid (TFA) and crosslinked peptides were enriched by peptide size exclusion chromatography (SuperdexPeptide 3.2/300 column, GE Healthcare) with a flow rate of 50 μl/ min in 30 % (v/v) acetonitrile with water and 0.1 % (v/v) TFA (Gomkale et al., 2021; Singh et al., 2020). Fractions of 50 μL were collected and early eluting fractions that contained crosslinked peptides were subjected to liquid chromatography-coupled -mass spectrometry (LC-MS)) analyses.

LC-MS analysis was performed as described (Gomkale et al., 2021; Singh et al., 2020). In brief, for each crosslinker crosslinked peptides were determined as technical duplicates using an Orbitrap QExactive HF Mass Spectrometer coupled to a Dionex UltiMate 3000 UHPLC system (both Thermo Fisher Scientific) that was equipped with an in house-packed C18 column (ReproSil-Pur 120 C18-AQ, 1.9 μm pore size, 75 μm inner diameter, 30 cm length, Dr. Maisch GmbH). First mass spectrometer (MS1) full scans were acquired with a resolution of 120,000, an injection time (IT) of 60 ms and an automatic gain control (AGC) target of 1 × 10^6^. Dynamic exclusion (DE) was set to 30 s and only charge states between 3 and 8 were considered for fragmentation. MS2 spectra were acquired of the 30 most abundant precursor ions; the resolution was set to 30,000; the IT to 128 ms and the AGC target to 1 × 10^5^. Fragmentation was enforced by higher-energy collisional dissociation (HCD) at 30% NCE.

Raw files were analyzed by pLink 2.3.9 (Chen et al., 2019) for the identification of crosslinked peptides against the full set of *H. sapiens* initiation factors and ribosomal proteins of the 40S subunit retrieved from the UniProt database (Bateman et al., 2021). False discovery rate (FDR) was set below 5% on spectrum level; crosslinked peptide spectrum matches (CSMs) were not evaluated manually. For each crosslinker identified crosslinks were first filtered to include only unambiguous crosslinks with ≥ 2 hits and a maximum -log_10_(E-value) ≥ 3 (from pLink 2.3.9) and then mapped onto our structural models of the open and closed h48S IC states using the software Xlink Analyzer 1.1.4 (Kosinski et al., 2015) and Chimera 1.15 (Pettersen et al., 2004). In total, we identified 748 and 714 unique crosslinks that could be mapped onto the models of the open and closed states, respectively. For the open state, 297 and 451 crosslinks could be mapped with LC-SDA and BS3, respectively, and for the closed state 280 crosslinks with LC-SDA and 437 with BS3.

---

**Figure 1 – figure supplement 1.**
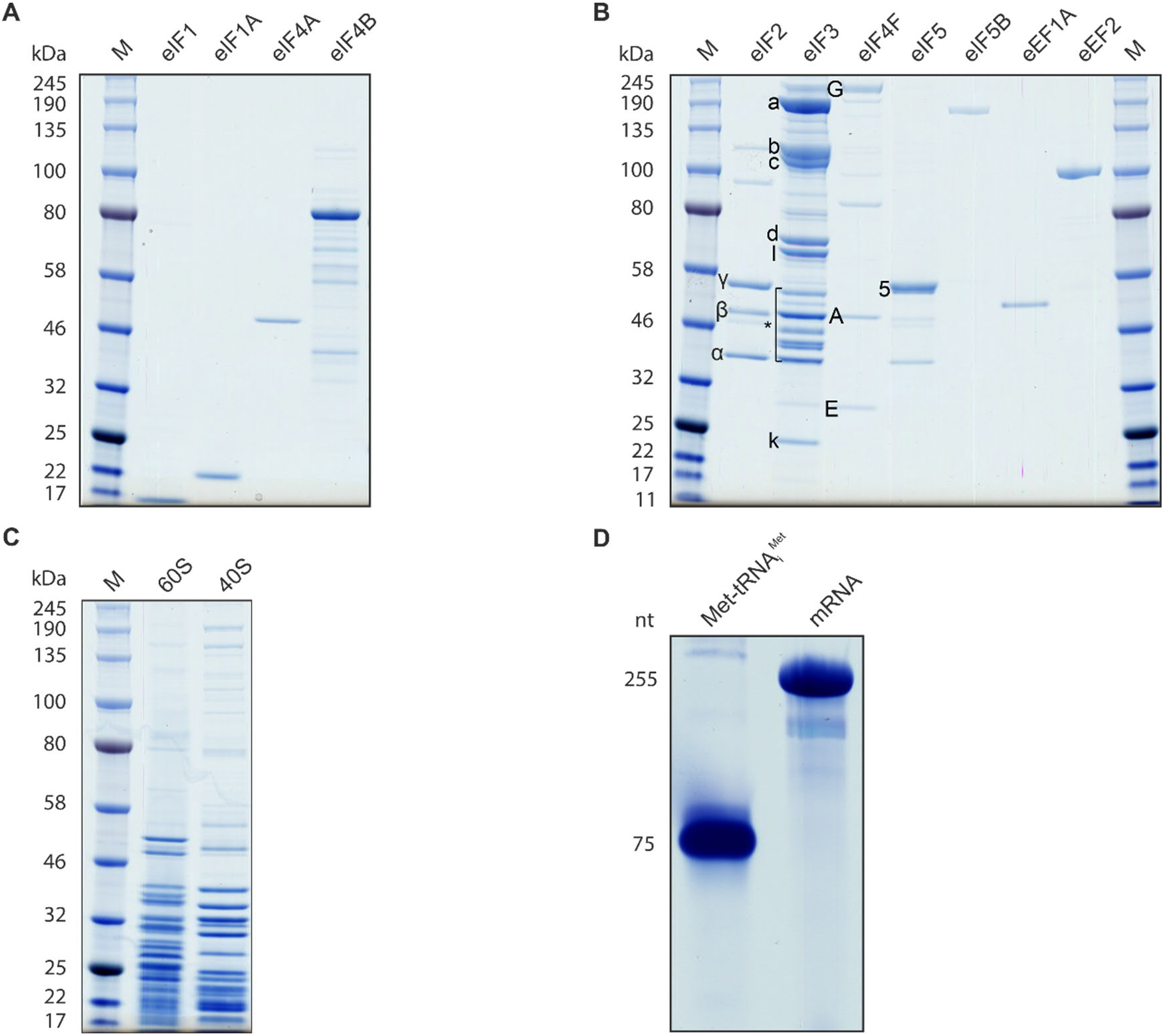
Components for the reconstituted *in vitro* translation system. **(A)** Recombinant human IFs: eIF1 (molecular weight 12.7 kDa), eIF1A (16.5 kDa), eIF4A (46.2 kDa) and eIF4B (69.2 kDa) expressed in *E. coli*. **(B)** Native initiation and elongation factors from mammalian cells. Proteins prepared from HeLa cells: eIF2 comprising 2α (36 kDa), 2β (38 kDa), and 2γ (52 kDa) subunits; eIF3 subunits 3a (167 kDa), 3b (92 kDa), 3c (105 kDa), 3d (64 kDa), 3e (52 kDa), 3f (38 kDa), 3g (36 kDa), 3h (40 kDa), 3i (36 kDa), 3j (29 kDa), 3k (25 kDa), 3l (67 kDa), 3m (43 kDa), *eIF3 subunits 3e to 3j are indicated in the bracket; eIF4F subunits 4G (175 kDa), 4A (46 kDa), 4E (25 kDa); eIF5B (138.8 kDa), and elongation factors, eEF1A (50 kDa) and eEF2 (95 kDa); eIF4F was not used in this work. Native eIF5 (49.2 kDa) was purified from rabbit reticulocyte lysate. **(C)** Proteins of 60S and 40S ribosomal subunits from HeLa cells. **(D)** *In vitro* transcribed human initiator tRNA_i_^Met^ and model mRNA. The tRNA_i_^Met^ was aminoacylated with [^3^H]-methionine and purified.

**Figure 1 – figure supplement 2.**
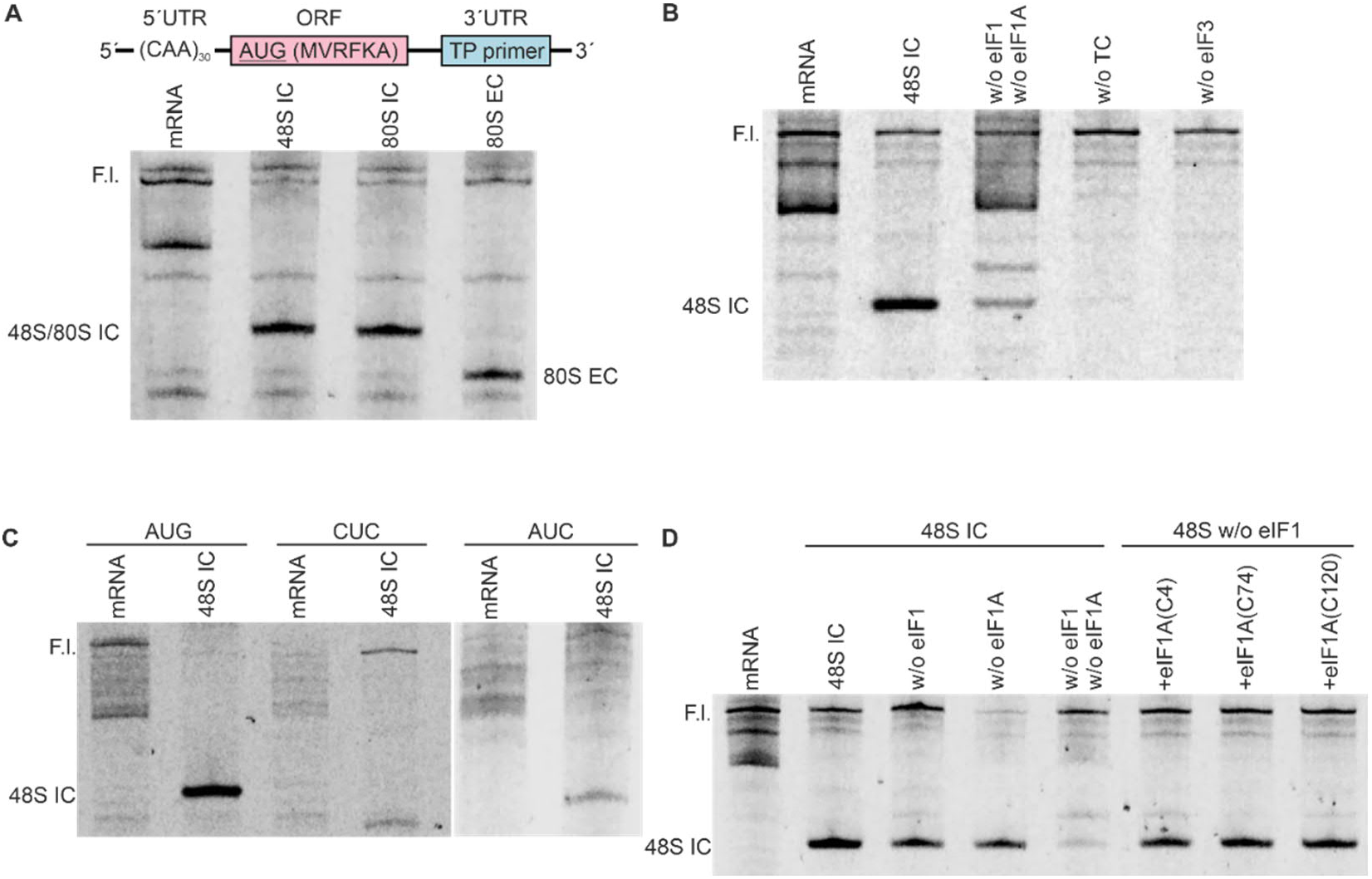
Fluorescence toe-printing assays of reconstituted h48S IC, h80S IC and h80S EC. **(A)** Schematic of the model mRNA and the toe-printing assay of the model mRNA without ribosome complex, and with h48S IC, h80S IC and h80S EC. The model mRNA is uncapped and has an unstructured 5’UTR containing 30 CAA repeats followed by an ORF coding for a short peptide MVRFKA and a toe-printing (TP) primer binding site in the 3’UTR. Toe-printing primer was labeled at the 5’ end by Atto647. The h48S IC was assembled from 40S subunits, eIF1, eIF1A, TC, eIF3, eIF4A and eIF4B in reaction buffer containing GTP. The h80S IC is assembled by adding eIF5, eIF5B and 60S subunits to h48S IC. 80S EC was formed upon synthesis of the MVRFKA peptide after addition of eEF1A, eEF2 with respective aminoacyl-tRNAs. F.l. is the full-length cDNA product of RT-reaction with the model mRNA as template. **(B)** Toe-printing assay of h48S IC in the absence (w/o) of individual eIFs. **(C)** Toe-printing of h48S IC assembled on AUG, CUC and AUC mRNA. No toe-print was detected for 48S CUC PIC and a weak cDNA band was found for 48S AUC PIC. **(D)** Toe-printing assay of h48S IC formed with eIF1A derivatives. The WT eIF1A was replaced by eIF1A derivatives with a cysteine at position 4, 74, or 120. Because the presence of either eIF1 or eIF1A alone can promote h48S IC formation on the model mRNA, the activity of eIF1A derivatives was tested in the absence of eIF1.

**Figure 1 – figure supplement 3.**
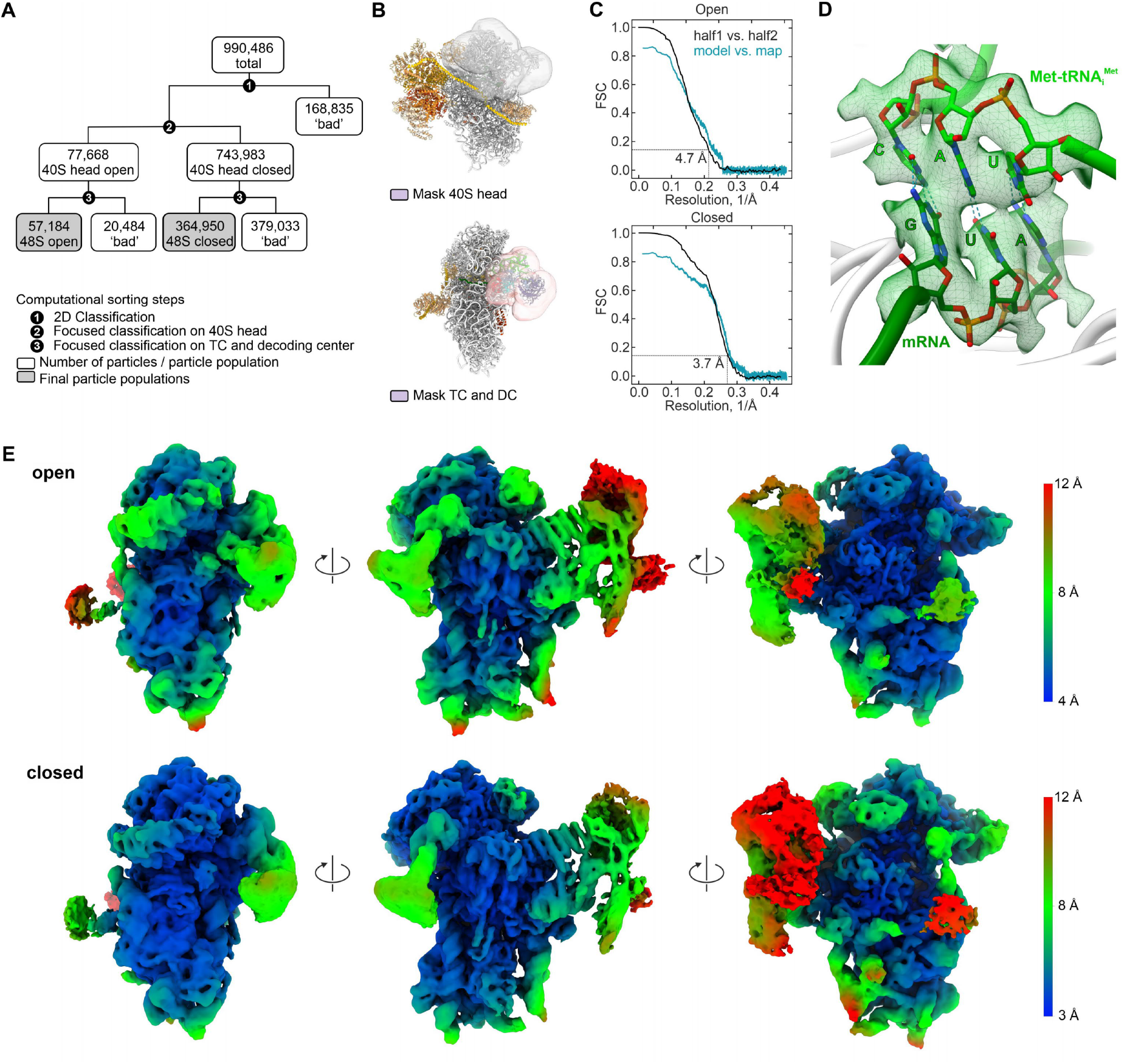
Cryo-EM data processing. **(A)** Sorting scheme for global and focused classifications of cryo-EM data. **(B)** Masks used for particle signal subtraction and focused classifications on 40S head (top) and ternary complex together with decoding center (bottom); DC, decoding center. **(C)** Fourier-Shell-Correlation (FSC) curves of the final cryo-EM reconstructions. **(D)** Details of the codon-anticodon interaction indicating start codon recognition in the closed state. The cryo-EM map (green transparent surface) is shown at 3σ. **(E)** Unsharpened cryo-EM maps of the open (top) and closed state (bottom) colored according to local resolution. Left: View onto TC and decoding center; center: View onto solvent site; right: View onto eIF3 core. Note the substantially lower local resolutions of eIF2γ and eIF3; atomic models for such regions were refined at lower resolution (see Methods).

**Figure 1 – figure supplement 4.**
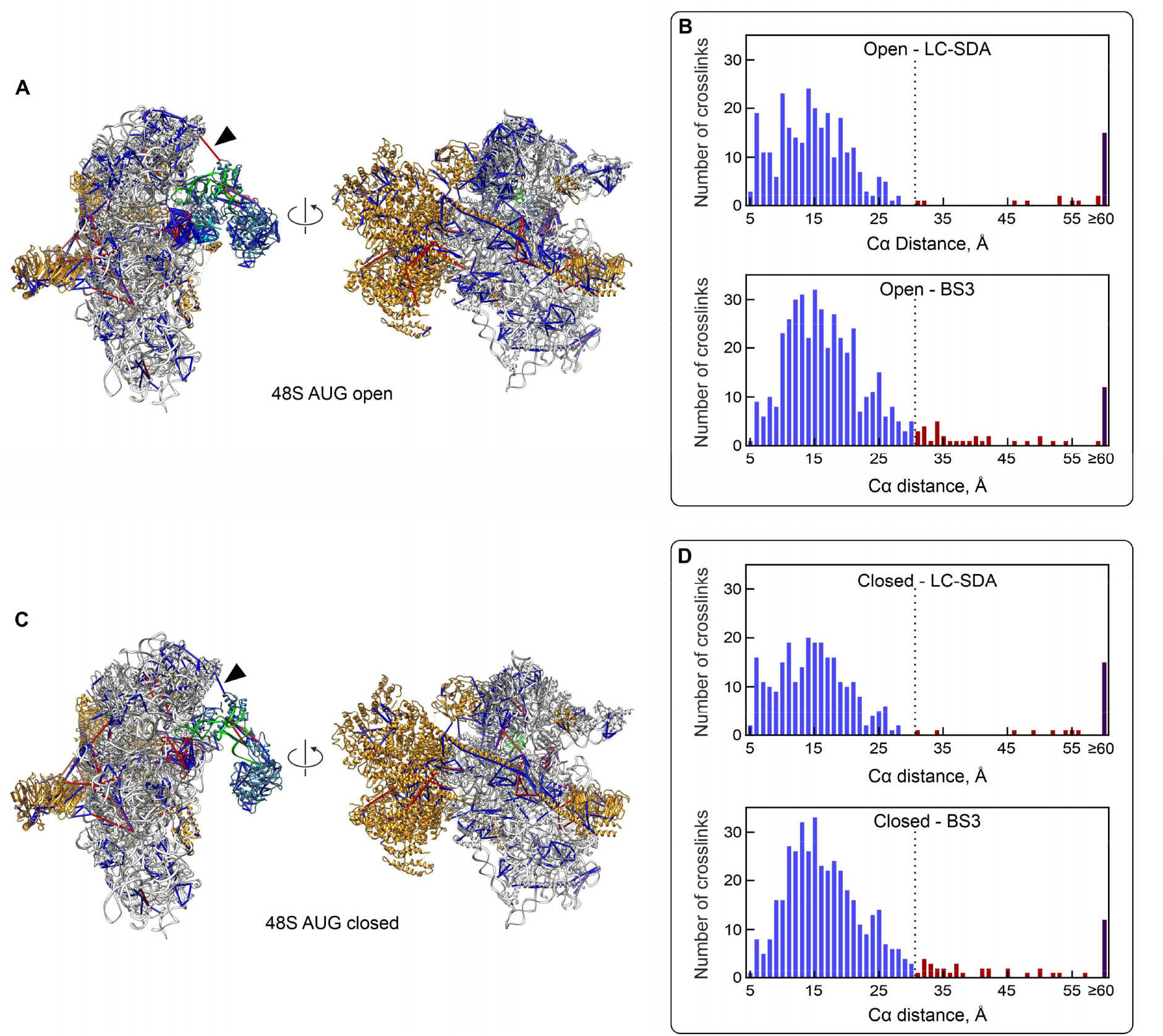
Crosslinking–mass spectrometry analysis. **(A)** and **(C)** Crosslinks mapped on the models of the open (A) and closed state (C), shown in two different orientations. Crosslinked residues are indicated by rods, colored in blue for permitted distances (<30 Å) and in red for non-permitted distances (30-45 Å); crosslinks for distances larger than 45 Å are not shown. The 40S subunit is depicted in white, Met-tRNA_i_^Met^ in green, eIF1 in cyan, eIF1A in red, eIF2 in steel blue and eIF3 in orange. The black arrow head marks the change in the crosslink distance between eIF2α (Lys123) and ribosomal protein S25 (Lys57) upon head closure. **(B)** and **(D)** Histograms of distances for unique crosslinks mapped onto the open (B) and closed states (D) obtained with the crosslinkers LC-SDA (top) and BS3 (bottom), respectively. Crosslinking distances longer than 45 Å account for approx. 6% of the crosslinks and most likely result from transient interactions of proteins not bound to the 40S subunit.

**Figure 2 – figure supplement 1.**
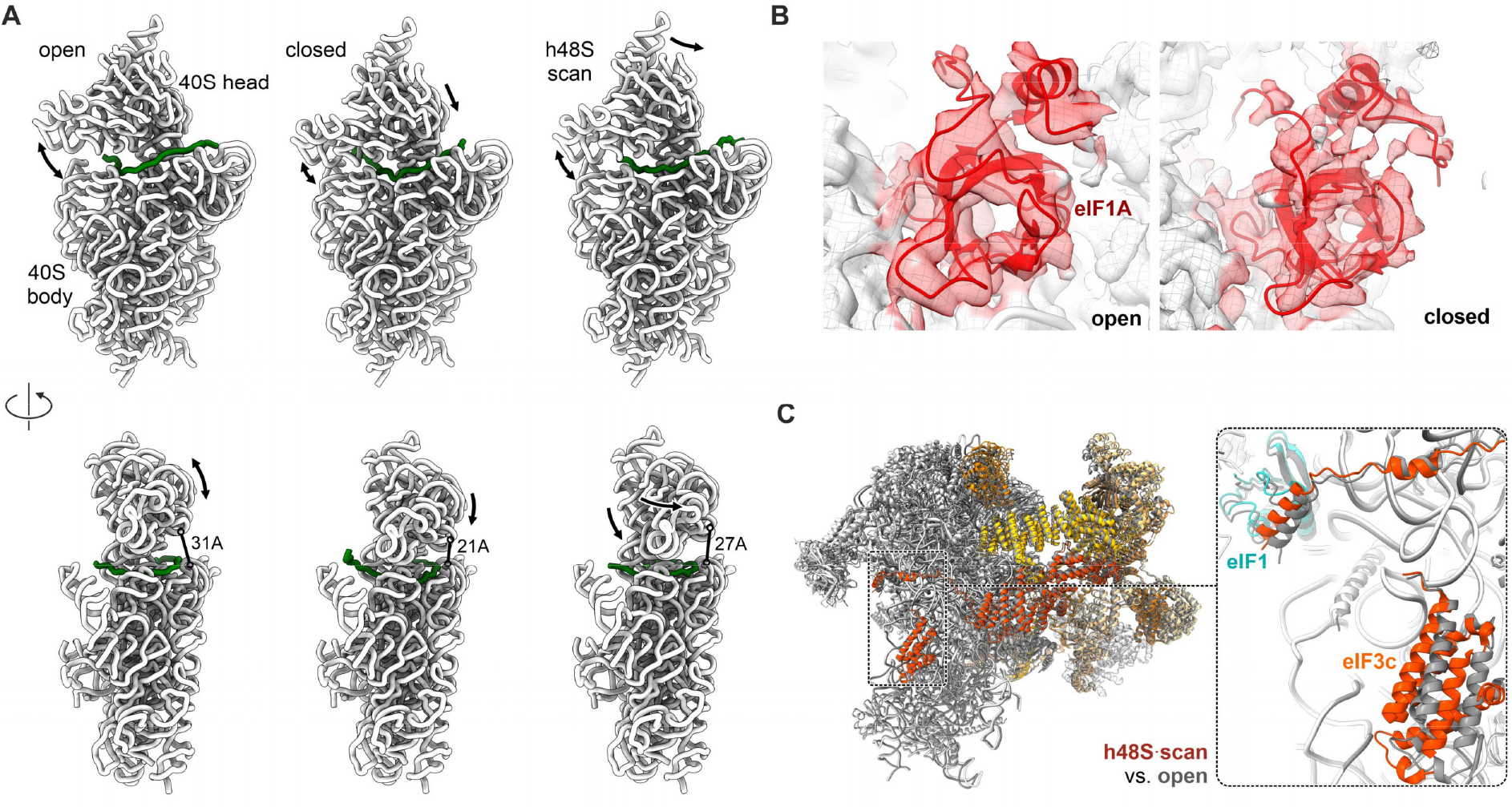
Structural rearrangements in h48S AUG IC. **(A)** Conformation of the 40S subunit in open, closed and scanning states. h48S scan, PDB 6ZMW (Brito Querido et al., 2020). For visual clarity, only the 18S rRNA (white) is shown with the mRNA (green). Arrows denote major modes of motion. Distances are measured between residues A1058 and A1640 of 18S rRNA. **(B)** Cryo-EM map of eIF1A in the open vs. closed state. The experimental map (transparent) is shown at 3σ with the ribbon models of eIF1A (red) and 18S rRNA (white). **(C)** eIF3 conformation. Left: Superposition of open h48S AUG (light grey) with h48S scan (white, eIF3 subunits in colors, PDB 6ZMW (Brito Querido et al., 2020)) revealing a similar overall eIF3 structure. Right: close-up showing a distinct orientation of the α-helix of eIF3c, which moves with eIF1, and a slight shift of its NTD between the two states.

**Figure 3 – figure supplement 1.**
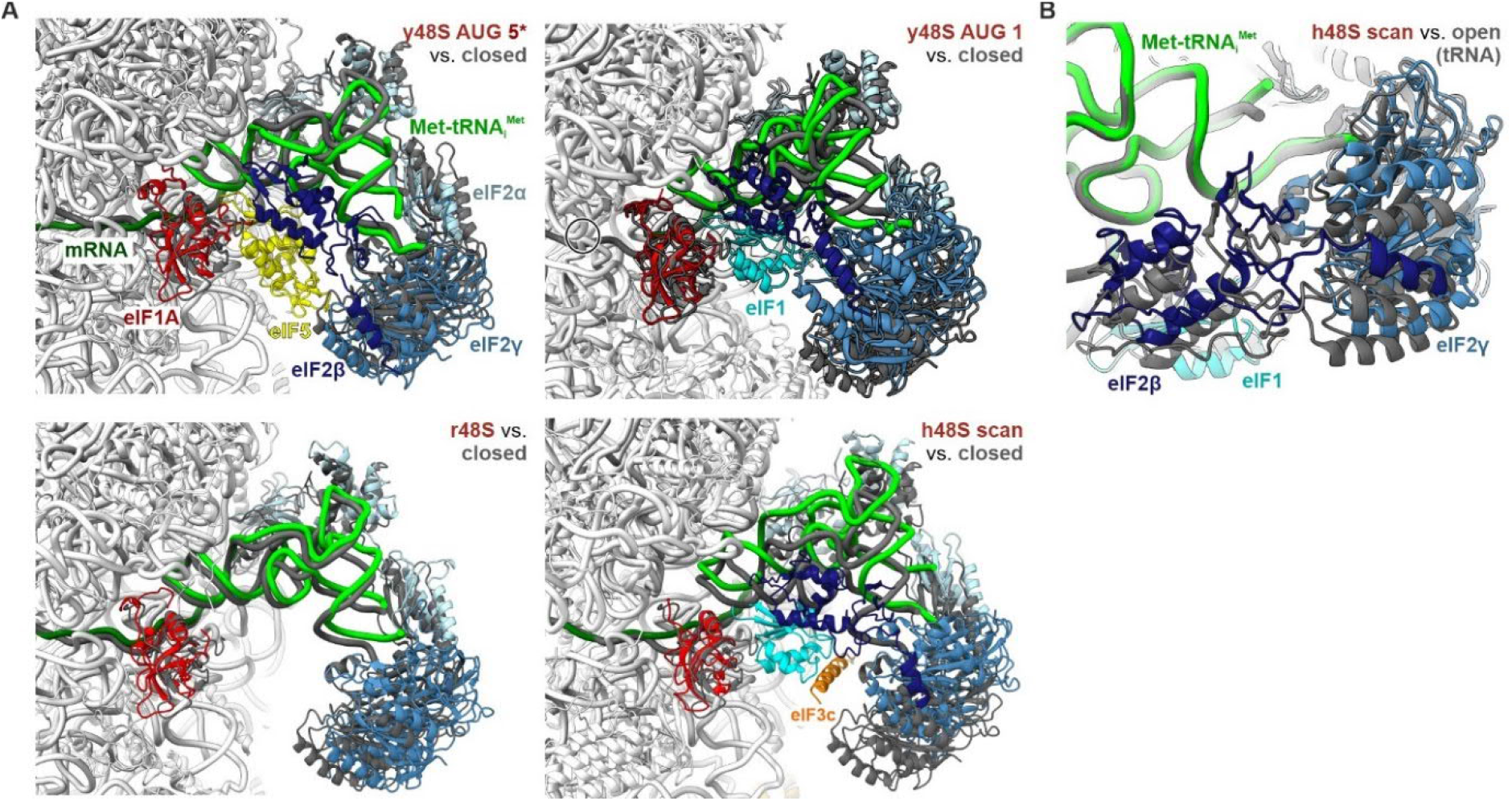
Comparison of the decoding center and TC in 48S ICs. **(A)** Superposition of the h48S AUG structures (light grey, 40S omitted) with other reported 48S ICs. y48S AUG 5*, cognate yeast 48S IC with eIF5 bound in state C2 (PDB 6FYY (Llacer et al., 2018)); y48S AUG 1, cognate yeast 48S IC with eIF1 bound in state C1 (PDB 4JAP (Llacer et al., 2015)); r48S AUG, rabbit 48S IC with cognate β-globin mRNA (Simonetti et al., 2020); h48S scan, non-cognate human 48S IC (PDB 6ZMW, (Brito Querido et al., 2020)). **(B)** Close-up of eIF2β and tRNA body. Alignment of h48S AUG open vs. h48S scan (PDB 6ZMW, (Brito Querido et al., 2020)).

**Figure 4 – figure supplement 1.**
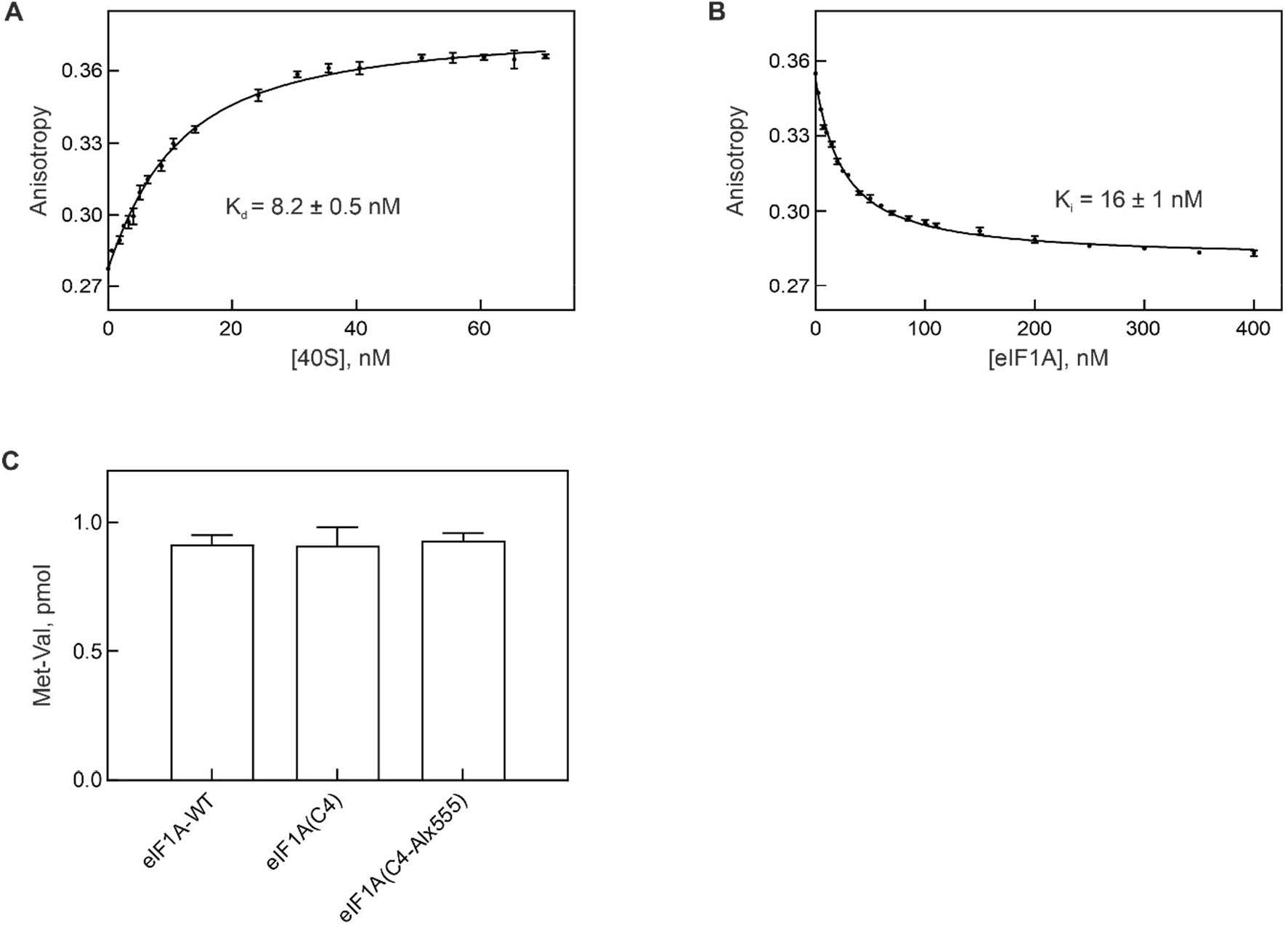
Functional activity of eIF1A(C4-Alx555). **(A)** Affinity of eIF1A(C4-Alx555) binding to the 40S subunit. eIF1A(C4-Alx555) (5 nM) was titrated with increasing concentrations of the 40S subunit at 25 °C. Anisotropy of eIF1A(C4-Alx555) was measured in a spectrofluorometer with excitation at 555 nm and emission at 568 nm. The binding curve was evaluated using a quadratic equation to obtain the binding affinity (K_d_). **(B)** Comparison of the binding affinities of fluorescence-labeled and unlabeled eIF1A using a competition titration assay. The 40S–eIF1A complex (5 nM) was titrated with increasing concentrations of unlabeled eIF1A. The IC_50_ of the inhibition curve and the K_i_ value (which reflects the K_d_ for unlabeled eIF1A) were calculated using GraphPad prism software. **(C)** Met-Val dipeptide formation on 80S EC assembled using eIF1A(wt), eIF1A(C4) and eIF1A(C4-Alx555). 80S IC (2 pmol) was assembled with the respective eIF1A variant and mixed with a 10-fold excess of eEF1A–GTP–[^14^C]Val-tRNA^Val^. The Met-Val dipeptide was analyzed by HPLC and quantified using [^14^C] radioactivity counting in the Met-Val and Val peaks.

**Figure 4 – figure supplement 2.**
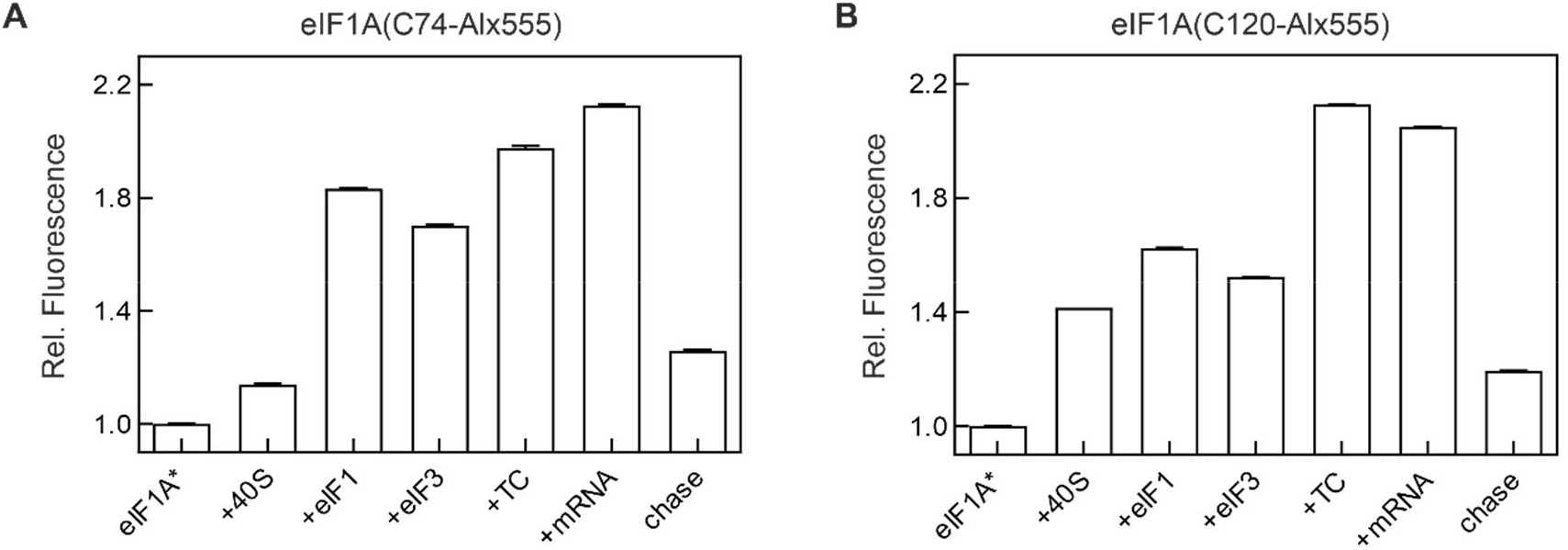
Fluorescence intensity changes of eIF1A(C74-Alx555) and eIF1A(C120-Alx555) upon 48S IC formation. **(A)** eIF1A(C74-Alx555). **(B)** eIF1A(C120-Alx555). Components of 48S IC were added sequentially in the order as indicated. Fluorescence intensity of free labeled eIF1A was set to 1.0. Error bars show standard deviation of 5 technical replicates (N=5).

## Author contributions

S.-H.Y., N.F., A.G, A.C., H.S., S.A., and M.V.R. conceived the experiments; S.-H.Y. and A.G. performed biochemical and kinetic experiments with the input of S.A.; S.-H.Y. analyzed the kinetic data; V.P., J.E.S., and N.F. performed cryo-EM experiments; A.L. and H.U. performed mass spectrometry analysis. S.-H.Y., V.P., N.F., S.A., and M.V.R. wrote the manuscript with input from all authors. All authors discussed the data analysis, critically reviewed the manuscript and approved the final version.

## Acknowledgements

We thank Prof. Dr. Tatyana Pestova for introducing us into the mammalian initiation system, generous guidance on biochemical material preparations and insightful comments on the manuscript; Thomas Conrad and Hossein Kohansal for HeLa cell extract; Olaf Geintzer, Vanessa Herold, Tessa Hübner, Franziska Hummel, Sandra Kappler, Christina Kothe, Anna Pfeifer, Lena Preiser, Theresia Steiger, and Michael Zimmermann for expert technical assistance. The work was funded by the Max Planck Society and the grants of the Deutsche Forschungsgemeinschaft (SFB860 to S.A., M.V.R. and H.S.; Leibniz Prize to M.V.R.). A.G. acknowledges the support of Boehringer Ingelheim Fonds.

## Declaration of interests

The authors declare no competing interests.

## Data availability

Final cryo-EM maps/atomic coordinates have been deposited in the EM database/Protein Data Bank with accession codes EMD-XXXX/PDB YYYY for the h48S AUG open state and EMD-XXXX/PDB YYYY for the h48S AUG closed state, respectively. Cryo-EM raw data have been deposited in the EMPIAR database as entry EMPIAR-ZZZZ.

## Notes

### Competing Interest Statement

The authors have declared no competing interest.

